# Stepwise Neofunctionalization of the NF-κB Family Member c-Rel during Vertebrate Evolution

**DOI:** 10.1101/2024.01.23.575293

**Authors:** Allison E. Daly, Abraham B. Chang, Prabhat Purbey, Kevin J. Williams, George Yeh, Shuxing Li, Scott D. Pope, Byrappa Venkatesh, Benjamin D. Redelings, Sibon Li, Kaylin Nguyen, Joseph Rodrigues, Kelsey Jorgensen, Trevor Siggers, Lin Chen, Stephen T. Smale

**Author notes:** These authors contributed equally (Allison E. Daly, Abraham B. Chang, Prabhat Purbey, Kevin J. Williams).

## Abstract

Adaptive immunity and the five vertebrate NF-κB/Rel family members first appeared in cartilaginous fish, suggesting that divergence and specialization within the NF-κB family helped facilitate the evolution of adaptive immunity. One specialized function of the NF-κB c-Rel protein in macrophages is the activation of *Il12b*, which encodes a key regulator of T-cell development. We found that c-Rel is a far more potent regulator of *Il12b* than of any other inducible genes in macrophages, with c-Rel regulation of *Il12b* dependent on its heightened intrinsic DNA-binding affinity. c-Rel homodimers regulate *Il12b* transcription in part via motifs with little resemblance to canonical NF-κB motifs. ChIP-seq experiments further defined distinct c-Rel DNA-binding preferences genome-wide, and X-ray crystallography of a c-Rel/RelA chimeric protein identified key amino acid changes that support the unique c-Rel properties. Unexpectedly, these changes, along with the c-Rel/RelA binding affinity differences, were largely restricted to mammalian species. Together, our findings reveal how a transcription factor family member can undergo a structural transition at a late stage of vertebrate evolution, resulting in an increased intrinsic DNA binding affinity and with clear functional consequences, presumably to support the increasing complexity of immune regulation.

## INTRODUCTION

The V(D)J adaptive immune system reflects billions of years of evolution and is known to rely on complex interactions between hundreds of distinct cell types. Its defining features, including immunoglobulin, T-cell receptor, and major histocompatibility complex genes, are apparent in cartilaginous fish that emerged approximately 450 million years ago and have been retained in all jawed vertebrates (Gnathostomata) that have been studied. In contrast, jawless fish (Cyclostomata), including lampreys and hagfish, which evolved from a common ancestor of gnathostomes, rely on variable lymphocyte receptors for antigen recognition.^1–3^

Although the V(D)J adaptive immune system is fundamentally similar among gnathostomes, differences have been observed that are likely to reflect increasing complexity to accommodate the needs of distinct vertebrate groups. For example, canonical immunoglobulin class switching first appeared in amphibians, and lymph nodes and germinal centers emerged in mammals.^3–5^ Additional changes in adaptive immune regulation were required to support fetal tolerance in placental mammals,^6^ as exemplified by a placental mammal-specific enhancer in the locus encoding FoxP3, a key transcription factor controlling Treg development.^7^ In addition to this enhancer, which supports extrathymic Treg development, functionally impactful amino acid changes in the FoxP3 protein occurred during distinct stages of vertebrate evolution.^8^

Multiple mechanisms by which gene duplication can support evolution have been described.^9–10^ Subfunctionalization occurs when the functions of an ancestral protein are divided between its duplicated descendants. Neofunctionalization represents the emergence of new functions in one or more descendant. Neofunctionalization often arises from non-coding mutations that alter the expression patterns of the duplicated genes, but mutations resulting in amino acid changes also allow new functions to emerge.^11–13^

The mammalian NF-κB/Rel transcription factor family is comprised of five members that play prominent roles in the regulation of both innate and adaptive immune responses.^14–15^ Invertebrate NF-κB ancestors possess well-documented roles in development, homeostasis, and innate immunity.^16^ NF-κB proteins in both invertebrates and vertebrates are characterized by an N-terminal Rel homology region (RHR) of approximately 300 amino acids that supports sequence-specific DNA binding and assembly into homodimeric and heterodimeric species.^15^ Most dimers are retained in the cytoplasm of resting cells in association with IκB inhibitor proteins, with IκB degradation and the nuclear translocation of the NF-κB dimers directed by signaling pathways upon sensing of microbial and environmental threats. Loss-of-function studies have revealed distinct biological functions for each NF-κB family member,^15,17,18^ but much less is known about the target genes and regulatory mechanisms for each dimeric species.

RelA and c-Rel are the two most closely related NF-κB family members in mammals, with highly conserved RHRs and identical DNA-contacting residues, but divergent C-terminal activation domains that are poorly conserved through evolution. RelA is abundant in most cell types, whereas high c-Rel expression is found primarily in hematopoietic lineages.^19^ Mice deficient in the *Rela* gene (encoding RelA) exhibit embryonic lethality due to broad deficiencies in cell survival.^15^ In contrast, mice deficient in the *Rel* gene (encoding c-Rel) exhibit a variety of immune abnormalities, including diminished T- and B-cell responses, defective Treg development, and a prominent loss of both Th1 and Th17 immune responses.^15,19,20,21^

In immune cells that express both RelA and c-Rel, the two proteins often act redundantly to regulate inducible genes.^15,22^ Nevertheless, c-Rel has been reported to contribute in a non-redundant manner to the induction of several genes.^19^ We and others previously described a highly potent role for c-Rel in the activation of the *Il12b* gene in both mouse and human antigen presenting cells, including macrophages and some dendritic cell subsets.^23–26^ The regulation of *Il12b* by c-Rel, which extends to patients with inherited human c-Rel deficiency,^26^ helps explain c-Rel’s established importance for Th1 and Th17 immune responses, as *Il12b* encodes a common subunit of the heterodimeric IL-12 and IL-23 cytokines needed for Th1 and Th17 cell development, respectively.^27^

In a chimeric protein analysis, the c-Rel requirement for *Il12b* induction localized to an 86-residue portion of the RHR containing 46 amino acid differences between RelA and c-Rel.^28^ Despite identical DNA-contacting residues in RelA and c-Rel,^29–30^ this 46-residue region allow c-Rel homodimers to bind NF-κB motifs with a much higher affinity than RelA homodimers.^28,31^ The finding that the same small region of c-Rel required for *Il12b* gene induction in *Rel^-/-^* macrophages also confers a large DNA affinity difference suggests that the affinity difference is responsible for the functional difference.

In this study, we examined the unique properties of c-Rel and their relevance for the functional specialization of c-Rel during vertebrate evolution. We first found that, in activated macrophages, *Il12b*’s strong dependence on c-Rel differs dramatically from the limited c-Rel dependence of almost all other inducible genes, including genes involved in innate immunity. We then found that c-Rel homodimers can bind motifs in vitro and in vivo that bear little resemblance to canonical NF-κB motifs due to the uniquely high intrinsic DNA-binding affinity of c-Rel homodimers. Importantly, the c-Rel-RelA affinity difference was not observed with DNA-binding domains from most non-mammalian species. X-ray crystallographic analysis of a mouse c-Rel-RelA chimeric protein revealed amino acids likely to explain the affinity difference, with the most important amino acid diverging between c-Rel and RelA only in mammalian species. Together, our results document a major structural transition within a transcription factor family member at a late stage of vertebrate evolution, presumably for the purpose of increasing functional divergence of family members and supporting the increasing complexity of adaptive immune regulation.

## RESULTS

### Evolution of the NF-κB/Rel Family

A full understanding of the NF-κB/Rel family of transcription factors will require an understanding of the family’s evolution. As an initial step toward this goal, we prepared a phylogenetic tree based on predicted amino acid sequences of the RHRs of NF-κB family members from several eukaryotic species (Figure 1). This analysis revealed that the five NF-κB family members - p50 (*Nfkb1*), p52 (*Nfkb2*), RelA (*Rela*), c-Rel (*Rel*), and RelB (*Relb*) – are conserved in all gnathostome species examined, including elephant shark, which is representative of cartilaginous fish (Chondrichthyes), the most primitive gnathostome class. Notably, the elephant shark genome encodes a sixth NF-κB protein that exhibits the closest homology to its p50 and p52 paralogs (Figure 1).

**Figure 1.**
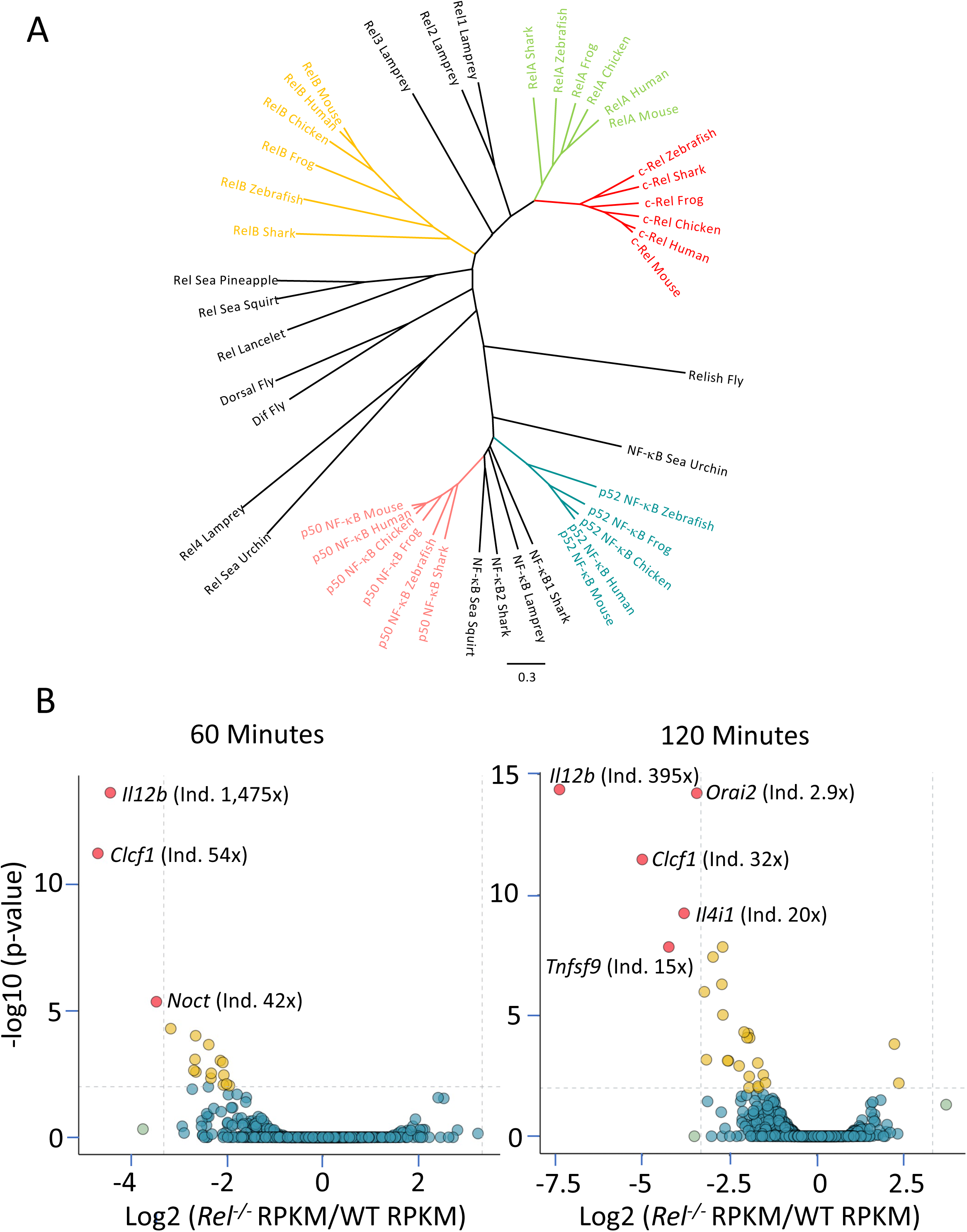
NF-κB Phylogenetic Analysis and Highly Selective c-Rel Requirement. (A) A phylogenetic tree was prepared with the RHR amino acid sequences of NF-κB family members from representative invertebrate and vertebrate species (see Methods). All sequences were from public databases. (B) Volcano plots were prepared comparing nascent transcript RNA-seq datasets from lipid A-induced genes in WT versus *Rel^-/-^* C56BL/6 macrophages. Comparisons are shown from cells stimulated with lipid A for 60 or 120 minutes. Genes exhibiting the strongest c-Rel dependence and weaker, statistically significant dependence are highlighted in red and gold, respectively. The identities of genes exhibiting the strongest c-Rel-dependence at each time point are included, along with their fold-induction in WT macrophages at that time point (in parenthesis).

Invertebrate NF-κB proteins (typically two or three in each species) diverge considerably from the vertebrate RelA, c-Rel, and RelB clusters (Figure 1). In contrast, a closer phylogenetic relationship is apparent between the vertebrate NF-κB p50 and p52 proteins and their invertebrate ancestors, including drosophila *Relish* (Figure 1A). Notably, the Relish precursor protein contains an ankyrin repeat domain, which is also characteristic of the p50 and p52 precursor proteins vertebrates.

Because the RelA, c-Rel, and RelB RHRs are evolutionarily distant from the invertebrate NF-κBs, it was of interest to examine NF-κB proteins in sea lamprey, a species representative of cyclostomes (jawless fish), which diverged from gnathostomes during the Ordovician period, roughly 450 million years ago.^2^ By mining a recently refined germline genome sequence for the sea lamprey, *Petromyzon marinus*,^32^ we identified five Rel/NF-κB genes, including three that were previously identified.^33^

One of the lamprey genes (Figure 1A NF-κB lamprey) clustered closely with two of the elephant shark p50/p52 proteins, and relatively closely with other vertebrate p50 and p52 proteins (Figure 1A, NF-κB lamprey). Like the p50 (*Nfkb1*) and p52 (*Nfkb2*) genes and the drosophila *Relish* gene, this gene encodes an ankyrin repeat domain (not shown). We speculate that a common cyclostomata/gnathostomata ancestral gene gave rise to this lamprey NF-κB gene and also, through gene duplication (presumably during a tetraploidization event),^34^ to the gnathostome p50 and p52 genes. However, none of the other four lamprey NF-κB proteins clustered closely with the RelA, c-Rel, and RelB proteins. These results suggest that the distinguishing features of RelA, c-Rel, and RelB first appeared after the cyclostomata/gnathostomata divergence.

A closer comparison of the RHR sequences from the four lamprey Rel proteins and the RelA and c-Rel proteins from various vertebrate species provides further support for the hypothesis that the lamprey RHRs lack the characteristic features of the RelA and c-Rel RHRs. Specifically, all four lamprey Rel RHRs contain a comparable number of mismatches (between 23 and 52) to the 159 RHR amino acids that are most highly conserved between both c-Rel and RelA in other vertebrates (Figure S1A, S1B). In addition, an analysis of 23 RHR residues that most consistently distinguish the RelA RHR from the c-Rel RHR revealed that none of the lamprey genes are strongly biased toward either the RelA or c-Rel distinguishing residues (Figure S1C). Because considerable evidence supports the existence of a genome tetraploidization event prior to cyclostome/gnathostome divergence, as well as additional tetraploidization events in both cyclostomes and gnathostomes after their divergence,^34,35^ the ancestral relationship between the four lamprey Rel proteins and the RelA and c-Rel proteins is difficult to predict. However, the fact that the c-Rel and RelA RHRs exhibit much greater similarity to each other than to any of the sea lamprey RHRs suggests that the c-Rel and RelA genes arose by duplication of a single common gnathostome ancestral gene. Moreover, because the c-Rel and RelA genes, like V(D)J adaptive immune systems, are characteristic of all gnathostomes, their divergence may have helped support the emergence of V(J)J adaptive immunity.

### Highly Selective Role of c-Rel in Lipid A-Stimulated BMDMs

RelA and c-Rel contain the two most closely related RHRs among the five vertebrate NF-κB family members, including identical DNA-contacting residues.^29,30^ Despite considerable redundancy between the two proteins in some settings, they contribute unique functions for the regulation of adaptive immunity.^15,18,25,36^ To improve our understanding of c-Rel’s functions in an innate immune cell type, we performed RNA-seq with wild-type C57BL/6 and *Rel^-/-^* bone marrow-derived macrophages (BMDMs) stimulated with lipid A for 0, 60, and 120 min to identify c-Rel-dependent genes at a genome-wide scale. This analysis was done by nascent transcript (chromatin-associated) RNA-seq (Figure 1B), to allow a quantitative analysis of c-Rel’s impact on transcription (because mRNA levels can also be strongly influenced by mRNA stability).

Surprisingly, these analyses revealed strong c-Rel-dependent expression of only a small number of genes one and two hr post-stimulation, when direct targets are expected to first be induced (Figure 1B). *Il12b* exhibited unusually strong c-Rel-dependence at both time points, and *Il12b* was induced by lipid A much more potently than the other genes that exhibited strong c-Rel-dependence (Figure 1B). The other c-Rel-dependent genes include *Clcf1* (encoding cardiotrophin-like cytokine factor 1), *Orai2* (encoding a calcium channel component), *Il4i1* (encoding a secreted L-amino acid oxidase), *Noct* (encoding a member of the exonuclease-endonuclease-phosphatase superfamily of enzymes), and *Tnfsf9* (encoding a transmembrane cytokine of the tumor necrosis factor family) (Figure 1B, fold induction values in parenthesis). The functions of these genes in cells of the innate immune system remain poorly understood. Notably, strong c-Rel dependence was not observed at these early time points at any genes known to play central roles in innate immune responses.

The primary function of the IL-12b (IL-12 p40) protein encoded by the *Il12b* gene is to regulate adaptive immune responses by promoting the differentiation of naïve T cells into the Th1 and Th17 lineages. Moreover, the regulation of *Il12b* expression by c-Rel has been shown to be critical for the development of Th1 and Th17 cells.^26,37,38^ Thus, c-Rel’s highly selective role as a potent inducer of *Il12b* is consistent with the hypothesis that, during vertebrate evolution, c-Rel’s neofunctionalization allowed it to support adaptive immunity via its critical *Il12b* regulatory function in macrophages, along with its numerous well-documented functions in T and B cell subsets.^18,19^

### Specific Binding of c-Rel Homodimers to Highly Divergent DNA Recognition Motifs

Protein-binding microarrays (PBMs) previously revealed that c-Rel and RelA homodimers bind similar distributions of DNA recognition motifs (Figure 2B, each black dot represents a different motif),^31^ resulting in the same position weight matrix (PWM) consensus sequence (Figure 2A).^31^

**Figure 2.**
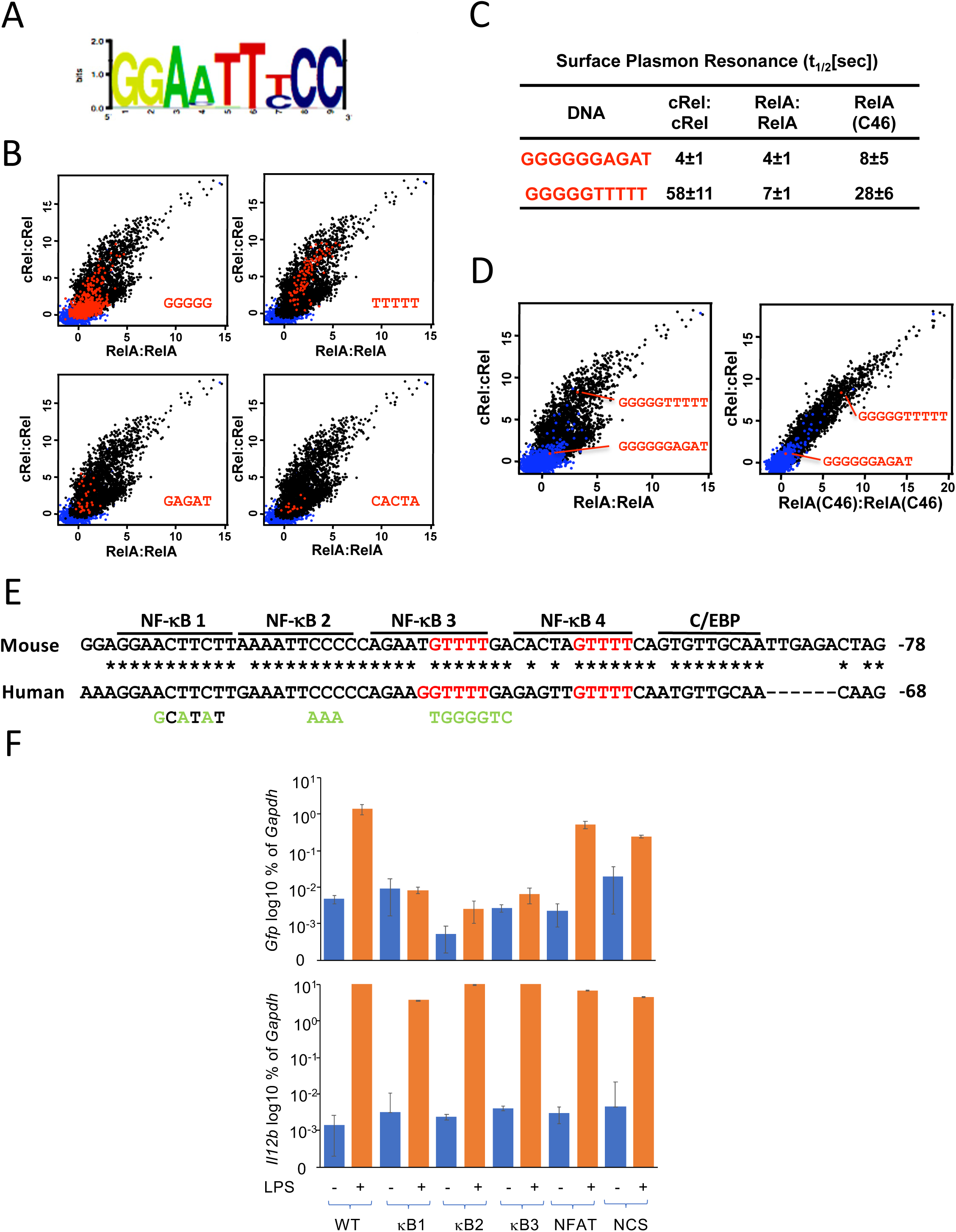
PBM and SPR Evidence of NF-κB Binding to Novel Motif Sequences. (A) A PWM-derived consensus recognition motif representative of the preferred binding sites for both RelA and c-Rel homodimers determined by PBM is shown.^31^ (B) PBM scatter plots are shown comparing the binding (in arbitration units) of recombinant RelA (x-axis) and c-Rel (y-axis) homodimers to a broad range of double-stranded oligonucleotide sequences. Oligonucleotides containing the sequence shown in each graph are highlighted in red. These graphs reveal binding to many oligonucleotides containing the sequence, TTTTT. By comparison, the proteins generally bind more weakly to oligonucleotides containing the sequences, GAGAT and CACTA, and to oligonucleotides containing, the sequence, GGGGG, despite the fact that RelA and c-Rel half-sites typically include two tandem G:C bps. The diagonal shape shows the RelA and c-Rel homodimers have similar spectrums of DNA preferences. However, the relative affinities of the two dimers for each motif cannot be determined from PBM data. (C) SPR analysis reveals that c-Rel homodimers bind a double-stranded oligonucleotide sequence, GGGGGTTTTT, with a much slower off-rate than the sequence, GGGGGGAGAT, further supporting a specific influence of the sequence, TTTTT, within one half-site. Off-rates are displayed as the time (seconds) needed for half of the protein to dissociate from the probe. RelA homodimers bind the sequence, GGGGGTTTTT, with a much faster off-rate than c-Rel homodimers, consistent with prior evidence that the intrinsic DNA-binding affinity of c-Rel homodimers (to all motifs) is much greater than that of RelA homodimers. The RelA (C46) chimeric protein binds with a much slower off-rate than RelA homodimers, confirming the importance of the 46 c-Rel residues for higher-affinity binding. (D) The locations of the oligonucleotide sequences in panel C are shown on PBM profiles comparing either RelA and c-Rel homodimers, or comparing c-Rel and RelA (C46) homodimers. (E) The sequences of portions of the mouse and human *Il12b* promoters are shown, with conserved nucleotides indicated (asterisks). The previously described non-consensus c-Rel homodimer binding sites (NF-κB1 and NF-κB2) and a previously described binding site for C/EBP proteins are indicated. Two additional conserved, potential NF-κB binding motifs (NF-κB3 and NF-κB4) are indicated. The GTTTT sequences that may help support NF-κB binding are highlighted in red. (F) A BAC encompassing the mouse *Il12b* locus was engineered in E. coli to contain *Gfp*-expressing sequences, and was then further engineered with substitution mutations in the *Il12b* NF-κB1, NF-κB2, or NF-κB3 sequences (Figure S2), with mutations introduced into a further downstream NFAT recognition sequence and a non-conserved sequence in the *Il12b* promoter as controls. The recombinant BACs were then stably integrated into mouse ESC and 2-6 independent clones containing single-copy BAC integrants were selected for each mutant. Following differentiation of the ESC into terminally differentiated macrophages, cells were stimulated with LPS for 0 or 2 hours, followed by mRNA isolation. qRT-PCR was then used to quantify relative mRNA levels in each clone for the endogenous *Il12b* mRNA (as a control confirming cell activation) and the BAC *Il12b-Gfp* mRNA (detected using *Gfp* primers). The graph displays mean relative mRNA levels and standard errors as a percentage of *Gapdh* mRNA level (y-axis, log10 scale) from 2-6 independent clones for the WT BAC clone and each BAC mutant.

However, surface plasmon resonance (SPR) experiments performed with representative oligonucleotides revealed that c-Rel homodimers bind with a much higher affinity than RelA homodimers to the full range of sequences evaluated by PBM.^31^ That is, the PBM results show relative binding of each individual protein to the thousands of DNA motifs present in the array, but SPR allows a comparison of the relative affinities of c-Rel homodimers and RelA homodimers for specific oligonucleotide sequences. The higher c-Rel homodimer binding affinity allows preferential c-Rel homodimer binding to motifs that diverge from the NF-κB consensus, as observed in early comparative studies.^39^

Interestingly, although most sequences bound by c-Rel and RelA homodimers in the PBM experiments show significant similarity to consensus NF-κB motifs, close scrutiny of the prior PBM profiles revealed that oligonucleotides characterized only by five tandem T:A bps were also bound by c-Rel and RelA homodimers (Figure 2B, top right, red dots). Many of these motifs lack the tandem G:C bps at the edges of the consensus motif (Figure 2A) that are considered to be the primary hallmarks of NF-κB consensus sequences and of NF-κB dimer binding. In contrast to the frequent binding to motifs with tandem T:A bps, binding was rarely observed with other DNA motifs that do not resemble known NF-κB recognition motifs (e.g. GAGAT and CACTA in Figure 2B) or a half-site motif that preferentially binds NF-κB p50 homodimers (Figure 2B, GGGGG).^31^

Frequent binding to oligonucleotides with tandem T:A bps was unexpected because structural and biochemical studies have revealed that NF-κB binding energy is primary due to contacts with G:C bps at the recognition sequence flanks (Figure 2A).^29^ However, SPR revealed that five T:A bps following an NF-κB half-site with five G:C bps results in a much slower c-Rel homodimer off-rate in comparison to an oligonucleotide in which the five G:C bps were followed by a different sequence (Figure 2C). This result suggests that tandem T:A bps are able to make a substantial contribution to c-Rel homodimer binding affinity. RelA homodimer binding to both oligonucleotides was much weaker by SPR (Figure 2C). The binding difference between c-Rel and RelA homodimers was largely due to the 46 key residues of c-Rel described above, as binding of the RelA(C46) chimeric protein to the motif with tandem T:A bps was readily detected by SPR, with an off-rate that was only moderately reduced in comparison to wild-type c-Rel (Figure 2C). Figure 2D shows the locations of the two motifs examined by SPR within the PBM profiles, adding further confirmation of the impact of tandem T:A bps on binding.

### Identification of Novel, Unrecognizable NF-κB Motifs in the *Il12b* Promoter

Unexpectedly, an examination of the mouse and human *Il12b* promoter sequences revealed that two conserved sequences with four tandem T:A bps each (Figure 2E, NF-κB3 and NF-κB4) are located immediately downstream of two previously described non-consensus NF-κB motifs (Figure 2E, *Il12b* NF-κB1 and NF-κB2).^28^ The *Il12b* NF-κB1 and NF-κB2 motifs, but not the *Il12b* NF-κB3 or NF-κB4 motifs, contain the two or three tandem G:C bp that can support stable binding in one half-site. Although the NF-κB3 and NF-κB4 motifs are unrecognizable as NF-κB motifs based on prior studies (due to the absence of tandem G:C bps), the data shown above, combined with conservation of the putative motifs in humans and mice (Figure 2E and below), led us to consider the possibility that they may contribute to c-Rel-dependent *Il12b* activation.

To examine the possibility of functional significance, we focused on the NF-κB3 motif, which was found to bind c-Rel dimers much more strongly than the NF-κB4 motif (see below). We first tested *Il12b* promoter-luciferase reporter plasmids containing the WT *Il12b* promoter and promoters with substitution mutations in the NF-κB1, NF-κB2, and NF-κB3 motifs. As previously shown,^40^ c-Rel overexpression in HEK 293 cells activated the WT *Il12b* promoter much more strongly than RelA overexpression (Figure S2). Mutations in each of the three NF-κB motifs reduced transactivation by c-Rel in this assay (Figure S2), supporting a possible role for the NF-κB3 motif.

To test the function of the NF-κB3 motif in a more native chromosomal context, we took advantage of a bacterial artificial chromosome (BAC) strategy developed prior to the emergence of CRISPR-Cas9 in mammalian cells (Figure S3; Doty et al., in preparation). Using a 200 kb BAC spanning the mouse *Il12b* locus, into which we had incorporated a green fluorescence protein cDNA (*Gfp*) to monitor BAC-derived *Il12b* expression, we introduced substitution mutations into the NF-κB1, NF-κB2, and NF-κB3 motifs (Figure 2E, bottom). Two additional mutations in the *Il12b* promoter (NFAT and NCS) were examined as controls. The WT and mutant *Il12b-Gfp* BACs were then stably transfected into mouse embryonic stem cells (ESCs; Figure S3A). After selection of multiple clones for each BAC containing single-copy BAC integrants, the clones were differentiated in vitro into terminally differentiated macrophages, followed by LPS stimulation for 2 hr (Figure S3A). qRT-PCR was used to monitor transcriptional induction of the *Il12b-Gfp* gene (using *Gfp* primers; Figure 2F, top). Transcription of the endogenous *Il12b* gene (using *Il12b* primers), which should be activated similarly in all WT and mutant lines, was also examined by qRT-PCR (Figure 2F, bottom). In this context, the WT *Il12b-Gfp* gene was activated, on average, greater than 100-fold (Figure 2F, top). In contrast, *Il12b-Gfp* activation was eliminated or greatly reduced in clones containing mutations in either the NF-κB1, NF-κB2, or NF-κB3 motif, whereas much smaller effects were observed with the two control mutations (Figure 2F, top). These findings further demonstrate important roles for all three NF-κB motifs.

### c-Rel Homodimer Binding to the Tandem *Il12b* Promoter Motifs

Electrophoretic mobility shift assay (EMSA) titration experiments with a recombinant protein containing the c-Rel RHR revealed that c-Rel homodimers can bind individually to each of the four non-consensus motifs from the *Il12b* promoter (Figure 3A), albeit with highly variable relative affinities, as demonstrated by quantification of EMSA complex intensities (Figure 3B). SPR experiments revealed the different c-Rel off-rates with the four motifs (Figure 3C), with NF-κB4 unable to bind with sufficient affinity for reliable results (data not shown). As expected, recombinant RelA homodimers bound poorly to all four sequences (Figure 3C). Notably, the chimeric RelA(C46) protein, which contains the 46 key residues of c-Rel in the context of the RelA protein, exhibited off-rates that were only moderately reduced in comparison to c-Rel (Figure 3C; see also Figure S4A for EMSA results with RelA(C46) protein).

**Figure 3.**
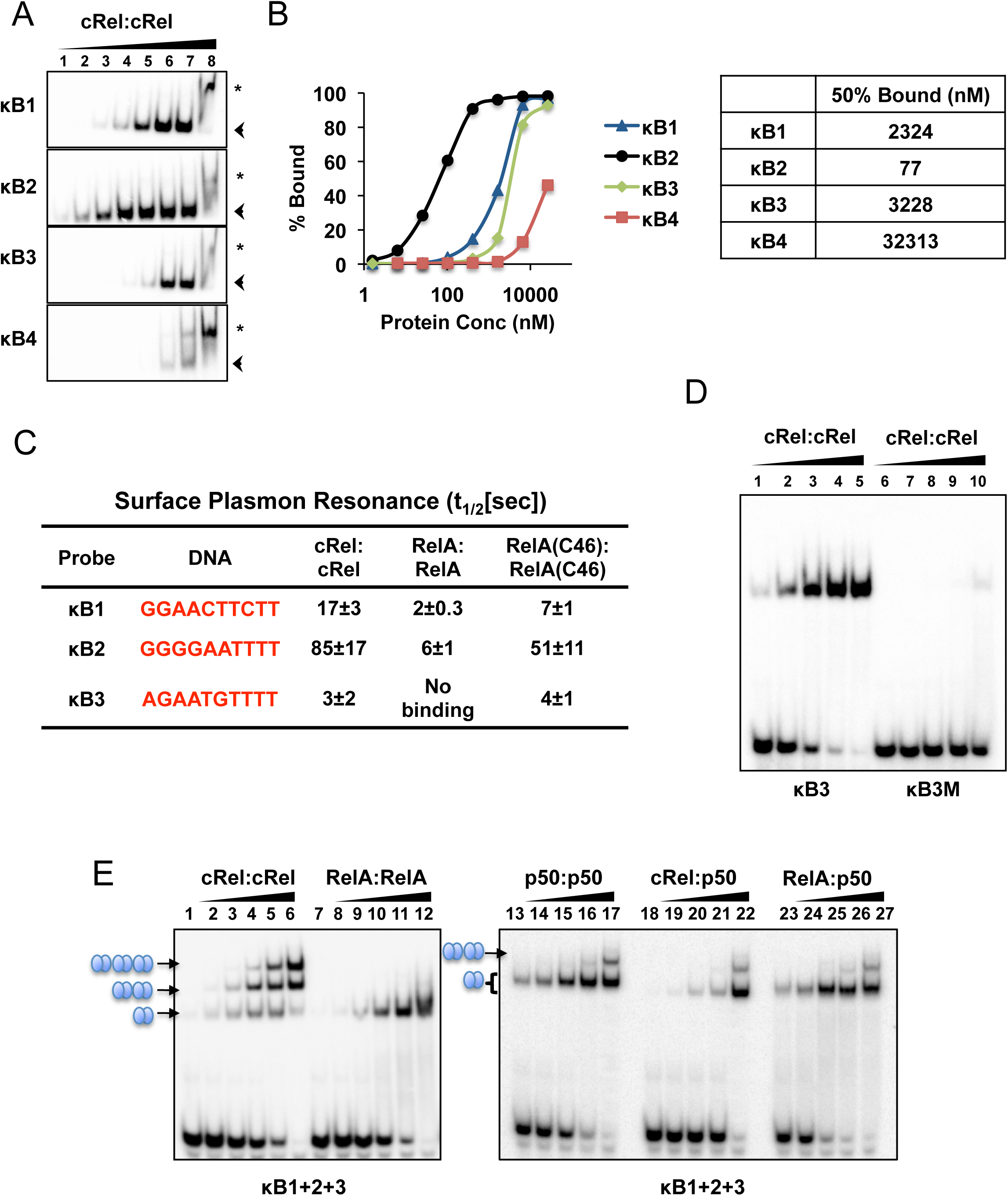
c-Rel Homodimer-Specific Binding to the *Il12b* Promoter Motifs. (A) EMSA experiments were performed with radiolabeled double-stranded oligonucleotides containing each of the four potential non-consensus NF-κB motifs from the mouse *Il12b* promoter. Increasing concentrations of recombinant c-Rel RHR homodimer protein expressed in E. coli were used. The locations of the predicted homodimer-DNA complexes are indicated (arrowheads), along with complexes that may represent aggregates formed at high protein concentrations (asterisks). (B) The line graph shows the percentage of each radiolabeled oligonucleotide probe from panel A bound by c-Rel protein at each protein concentration. The table (right) shows the concentration of c-Rel protein (nM) required to bind 50% of the radiolabeled probe. (C) SPR was used to determine off-rates of c-Rel, RelA, and RelA (C46) RHR homodimers from the *Il12b* NF-κB1, NF-κB2, and NF-κB3 motifs. No reproducible binding was observed with the NF-κB4 sequence. (D) EMSAs used to examine binding of increasing concentrations of the recombinant c-Rel RHR protein to radiolabeled, double-stranded oligonucleotide probes containing the WT NF-κB3 motif and a motif with multiple nucleotide substitutions (κB3M). (E) EMSAs were used to examine binding of purified recombinant c-Rel:c-Rel, RelA:RelA, p50:p50: c-Rel:p50, and RelA:p50 RHR dimers to a radiolabeled oligonucleotide probe containing the *Il12b* promoter sequence spanning the NF-κB1, NF-κB2, and NF-κB3 motifs. The locations of complexes containing one, two, or three bound dimers are indicated. The unbound radiolabeled probe is visible at the bottom of the image.

A comparison of six recombinant homodimeric and heterodimeric species - c-Rel:c-Rel, RelA:RelA, p50:p50, c-Rel:p50, RelA:p50 and RelB:p50 - revealed much stronger binding by c-Rel:c-Rel homodimers to both the NF-κB3 and NF-B4 motifs than by any of the other dimers (Figure S4B,C). In addition, an oligonucleotide containing substitution mutations in the tandem T:A bps of the NF-κB3 oligonucleotide exhibited greatly reduced binding in comparison to the WT NF-κB3 sequence, confirming the critical importance of the T:A bps (Figure 3D).

Notably, in EMSA titration experiments using a radiolabeled probe containing the native NF-κB1, NF-κB2, and NF-κB3 sequences from the *Il12b* promoter, efficient and simultaneous binding of three c-Rel homodimers was observed (Figure 3E). Mutation of any of the three motifs eliminated the slowest mobility band (Figure S5A). This property was unique to c-Rel homodimers, as only one RelA homodimer was capable of binding the same oligonucleotide, presumably due to the much lower affinity of this homodimer for each of the three motifs (Figure 3E). p50:p50, p50:c-Rel, and p50:RelA dimers were also incapable of loading three dimers onto this oligonucleotide (Figure 3E, right). Importantly, four c-Rel homodimers could simultaneously bind a radiolabeled probe containing all four potential recognition sequences from the *Il12b* promoter (Figure S5B).

A careful examination of the relative abundances of complexes containing one, two, three, or four bound dimers suggested the possibility of cooperative binding. To examine this possibility, we performed gel shift titrations with different combinations of two of the tandem motifs. Cooperative binding was not observed between the NF-κB1 and NF-κB2 motifs or between the NF-κB3 and NF-κB4 motifs (Figure S6). However, strong cooperativity between the NF-κB2 and NF-κB3 motifs was observed, in that the efficiency of c-Rel homodimer binding to NF-κB3 was greatly enhanced when the probe contained an adjacent intact NF-κB2 motif (Figure S6).

Together, these results strongly support a hypothesis in which preferential binding of c-Rel homodimers to three and possibly four tandem recognition motifs in the *Il12b* promoter underlies the c-Rel dependence of *Il12b* transcription. Moreover, the *Il12b* NF-κB3 and NF-κB4 motifs would have been entirely unrecognizable on the basis of past knowledge of NF-κB dimer binding specificities.

### c-Rel Versus RelA Preferential Binding In Vivo

To determine whether preferential binding of c-Rel can be observed in vivo, we performed ChIP-seq in BMDMs stimulated with lipid A for 0 and 60 min using c-Rel and RelA antibodies (three biological replicates with both antibody). To increase confidence in the subsequent analysis, we focused on the strongest 6,700 peaks obtained with each antibody, which upon integration of data sets yielded 8,134 distinct ChIP-seq peaks. We then determined their RPKM ratio at each of these 8,134 genomic locations (Figure 4A). This analysis revealed that the c-Rel/RelA binding ratio was comparable at most peaks, with 93% of peaks (7,549 of 8,134) exhibiting binding ratios between 1.3 and 3 (Figure 4A, right). However, stronger preferences for either c-Rel or RelA were observed at the remaining peaks. The median c-Rel/RelA RPKM ratio was 1.89, reflecting either differences in antibody quality or slightly stronger c-Rel interactions (or slightly more efficient c-Rel crosslinking) throughout the genome.

**Figure 4.**
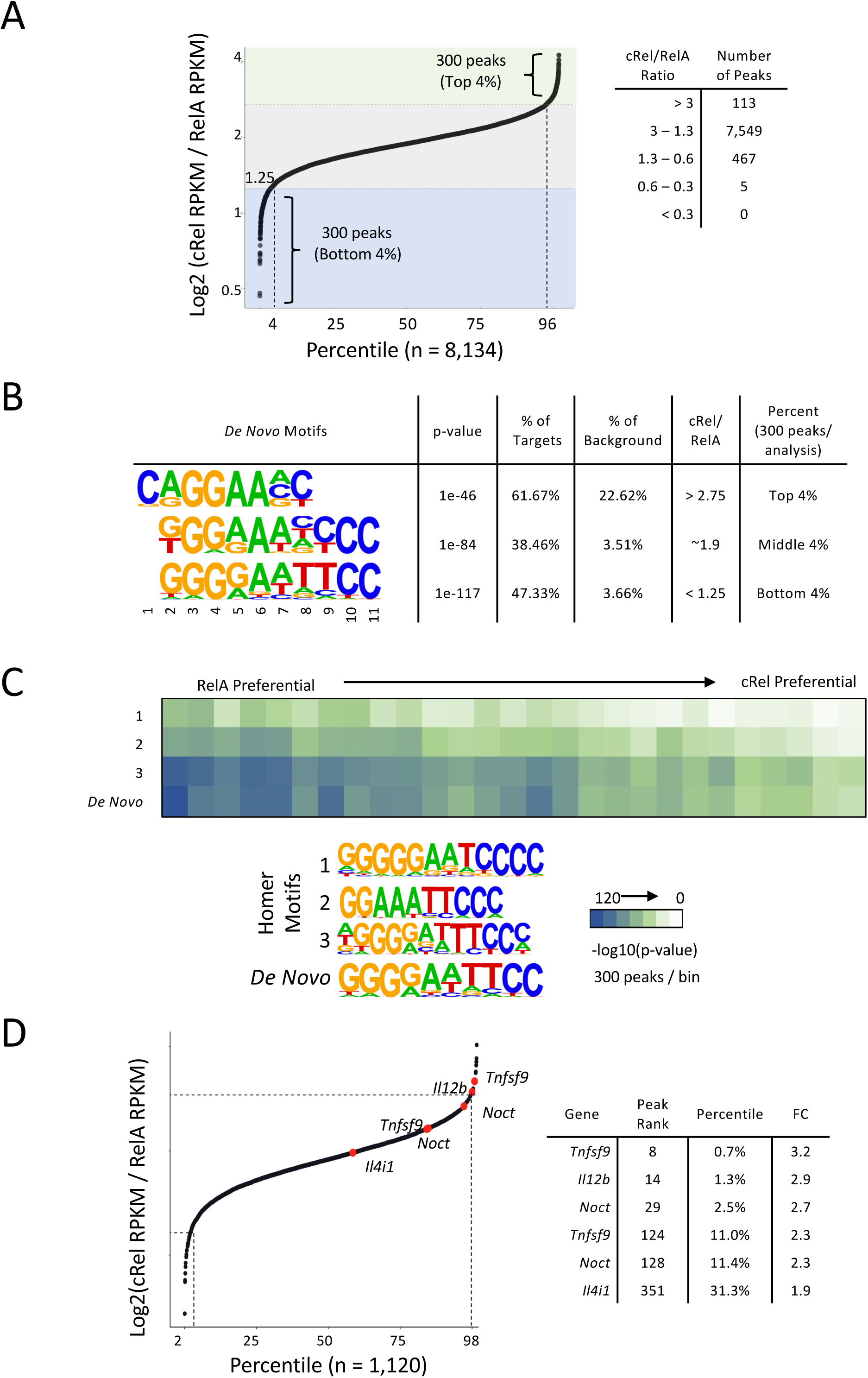
Selective Binding of c-Rel and RelA in Mouse BMDMs Examined by ChIP-seq. (A) ChIP-seq experiments were performed with antibodies to RelA and c-Rel in BMDMs stimulated with lipid A for 0 and 1 hour. The 6,700 peaks with the strongest peak scores obtained with each antibody at the 1-hour time point were selected and merged, yielding 8,134 peaks. c-Rel:RelA ratios were calculated from the RPKMs at each peak and were plotted (ratios on the y-axis), with the 8,134 peaks along the x-axis (displayed as a percentage of total peaks). The top and bottom 300 peaks based on ratio (4%) were then selected for motif analysis. At the right, numbers of peaks with different ranges of RPKM ratios are shown. (B) The most enriched motifs from a de novo motif analysis performed with Homer are shown. Motif analysis was performed with 300 peaks representing the largest peak ratios, 300 peaks representing the smallest peak ratios, and 300 peaks from the middle of the ratio distribution. The top motif identified by Homer with each set of peaks is shown. (C) The 8,134 peaks were divided into 27 bins of equal size across the spectrum of c-Rel:RelA RPKM ratios. The enrichment of three consensus NF-κB motifs within the Homer program, and the consensus NF-κB motif defined in panel B, was then examined across the spectrum. (D) Among the 8,134 peaks, 1,120 annotate to promoter regions (−1kb-+1kb relative to the TSS of annotated genes). The c-Rel/RelA RPKM ratios of these peaks were plotted and used to examine the relative ratios of peaks within the promoters of genes that exhibited strong c-Rel-dependence in Figure 1B. The *Il12b* promoter peak and one of two peaks within the *Tnfsf9* promoter ranked among the top 1.3% of peaks with the largest c-Rel:RelA RPKM ratios. Lesser c-Rel preferences were observed at a second peak in the *Tnfsf9* promoter, at two peaks that annotated to the *Noct* promoter, and at a peak in the *Il4i1* promoter. The ratio rank of each of these six peaks, the percentile among all promoter peaks, and the c-Rel:RelA RPKM ratio (FC) are shown at the right.

To determine whether selective binding by either c-Rel or RelA coincides with DNA sequence motif preferences, we performed de novo motif analyses with the 300 peaks exhibiting the largest and smallest RPKM ratios (top and bottom 4% of all peaks). Strikingly, this analysis revealed that the peaks exhibiting the largest c-Rel/RelA RPKM ratios (i.e. preferential c-Rel binding) showed the strongest enrichment for motifs resembling an NF-κB half-site (Figure 4B, GGAA) with two tandem G:C bps followed by two A:T bps. The enrichment of a half-site rather than a full dimeric NF-κB consensus motif (see Figure 2A) is consistent with the hypothesis that the enhanced binding affinity of c-Rel homodimers observed in vitro allows preferential c-Rel binding to many sites throughout the genome that diverge from consensus dimeric recognition motifs (i.e. motifs with two half-sites containing tandem G:C bps). In fact, a full consensus dimeric NF-κB motif was not among any of the four most enriched motifs exhibiting preferential c-Rel binding in this analysis (data not shown).

In contrast to the c-Rel-preferential peaks, RelA-preferential peaks exhibited strong enrichment of a motif resembling full consensus dimeric motifs defined in vitro for RelA:p50 or c-Rel:p50 heterodimers,^31^ with three tandem G:C bps at one half-site (optimal for p50 interactions), two tandem G:C bps at the other half-site (optimal for RelA or c-Rel interactions), and five intervening bps (Figure 4B). Enrichment of this conventional heterodimer motif suggests that the RelA interactions at sites exhibiting the lowest c-Rel/RelA binding ratios are mediated by RelA:p50 heterodimers. Because RelA:p50 heterodimers do not appear to bind DNA with a higher affinity than c-Rel:p50 heterodimers in vitro,^31^ this finding could reflect a high abundance of RelA:p50 heterodimers in comparison to c-Rel:p50 heterodimers or noise in the analysis. Alternatively, RelA:p50 interactions with a small subset of consensus motifs may be selectively stabilized in vivo by proteins that interact specifically with RelA.^41^

We also performed a de novo motif analysis with 300 peaks from the middle of the c-Rel/RelA RPKM ratio spectrum (Figure 4B). Here, we observed strong enrichment of a near-consensus dimeric NF-κB motif. The composition of this enriched motif suggests that it could represent interactions by a mixture of dimeric species.

We next examined how the enrichments of four distinct dimeric NF-κB consensus motifs (three motifs from the Homer program and one de novo motif from the above analysis) changes across the c-Rel/RelA RPKM ratio spectrum, after dividing the 8,134 peaks into 27 bins. This analysis revealed a consistent trend toward weaker enrichment of all four consensus motifs as the c-Rel/RelA RPKM ratio increases, again consistent with the view that c-Rel homodimers preferentially bind sequences that diverge from dimeric NF-κB consensus sequences.

Most importantly, we graphed c-Rel/RelA RPKM ratios for 1,120 peaks that annotate to promoter regions (−1kb to +1kb relative to the transcription start site [TSS]) (Figure 4D). This analysis revealed strongly preferential c-Rel binding to peaks within the promoters of three genes that exhibited strong c-Rel-dependent transcription, including the peak within the *Il12b* promoter, a peak in the *Tnfsf9* promoter region, and a peak in the *Noct* promoter, with binding that was less preferential at the *Il4i1* promoter and at additional independent peaks that annotate to the *Tnfsf9* and *Noct* promoters (Figure 4D). Thus, these ChIP-seq results add support to the structural and biochemical evidence of NF-κB dimer-specific binding preferences, with evidence that these binding preferences are relevant to the regulation of the *Il12b* gene and at least a few other c-Rel-dependent genes.

### c-Rel Versus p50 Preferential Binding In Vivo

We next used the same approach as above to examine preferential binding by c-Rel in comparison to p50. Because p50 ChIP-seq yielded a smaller number of peaks (probably due to lower antibody quality), this analysis was limited to the top 2,891 peaks obtained with each antibody, which, when merged, yielded 4,414 peaks (Figure 5A). Among these peaks, c-Rel ChIP-seq signals (RPKM) were generally stronger than p50 signals, with 2,646 peaks exhibiting c-Rel/p50 RPKM ratios greater than 3 (Figure 5A, right).

**Figure 5.**
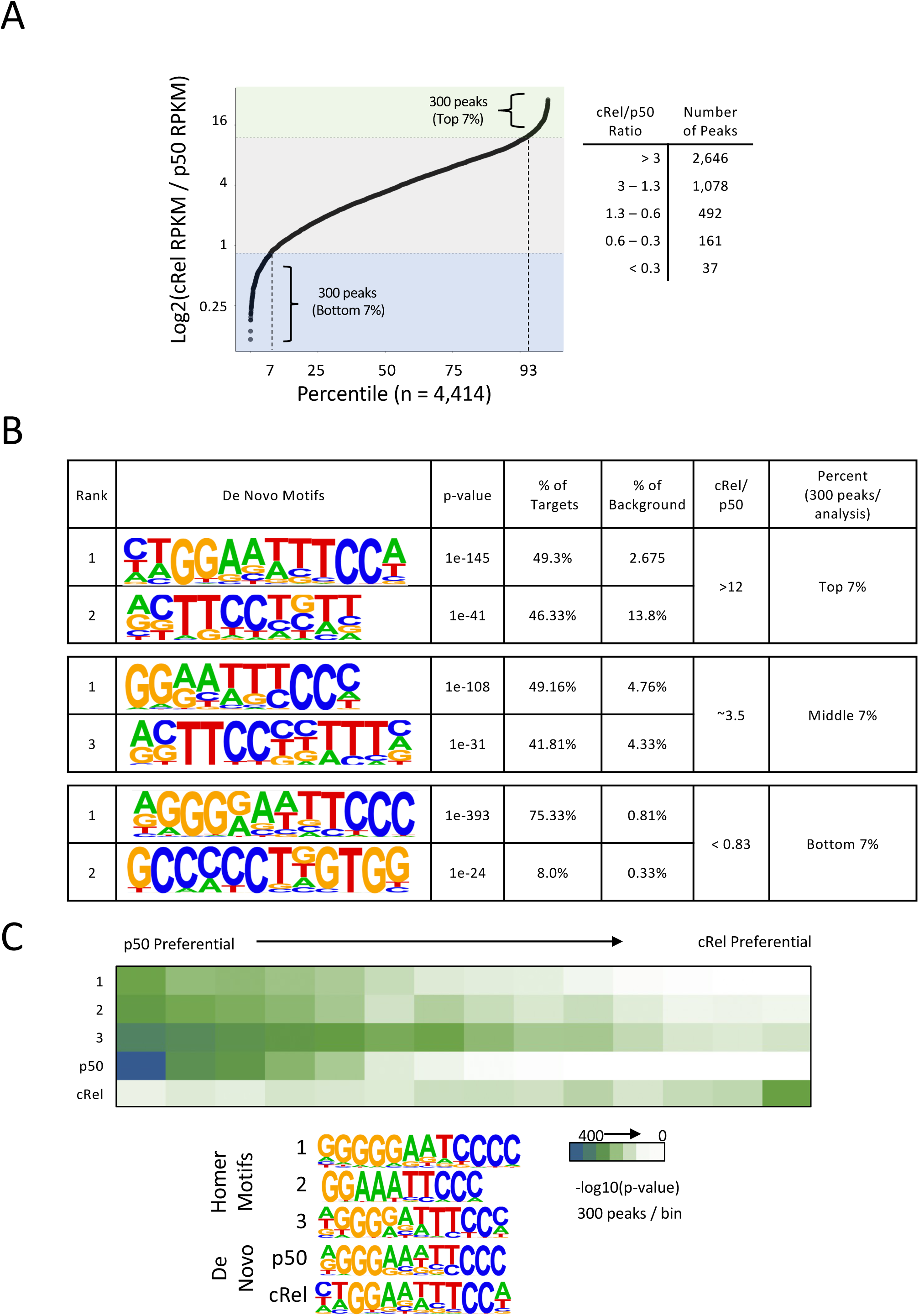
Selective Binding of c-Rel and p50 in Mouse BMDMs Examined by ChIP-seq. (A) ChIP-seq experiments were performed with antibodies to c-Rel and p50 in BMDMs stimulated with lipid A for 0 and 1 hourr. The 2,891 peaks with the strongest peak scores obtained with each antibody at the 1-hour time point were selected and merged, yielding 4,414 peaks (the peak number is smaller than in the RelA/c-Rel comparison because of the small number of peaks obtained with p50 antibody, probably due to relatively poor antibody quality). c-Rel:p50 RPKM ratios were then calculated at each peak and were plotted (ratios on the y-axis), with the 4,414 peaks along the x-axis (displayed as a percentage of total peaks). The top and bottom 300 peaks based on ratio (7%) were then selected for motif analysis. At the right, numbers of peaks with different ranges of RPKM ratios are shown. (B) The two most enriched motifs from de novo motif analyses performed with Homer are shown for three different groups of peaks. Motif analysis was performed with 300 peaks (7%) representing the largest peak ratios, 300 peaks representing the smallest peak ratios, and 300 peaks from the middle of the ratio distribution. (C) The 4,414 peaks were divided into 15 bins of equal size across the spectrum of c-Rel:p50 RPKM ratios. The enrichment of the three consensus NF-κB motifs within the Homer program, as well as one of the p50-preferential motifs and one of the c-Rel-preferential motifs identified in panel B, was then examined across the spectrum. NF-κB half-sites (not shown) were equally enriched across the spectrum, consistent with the presence of half-sites in all of the full consensus motifs examined.

De novo motif analysis performed with 300 peaks showing the largest c-Rel preference (top 7% of all peaks) revealed strong enrichment of two types of motifs (Figure 5B): The most enriched motif matched an anticipated consensus sequence for c-Rel homodimer binding, with two G:C bps within each half-site (Figure 5B, top). Strong enrichment was also observed with a motif characteristic of an NF-κB half-site, providing further evidence that c-Rel homodimers are unique in their preferential binding to sites that lack a dimeric consensus sequence (Figure 5B). In contrast, motifs showing the smallest c-Rel/p50 RPKM ratios exhibited characteristics of p50 homodimer binding. The most enriched motif contained three G:C bps within each half-site (separated by five bp), consistent with p50 interactions with three G:C bps (Figure 5B, bottom). Strong enrichment was also observed with a motif containing five tandem G:C bps, which perfectly matches a de novo motif that exhibited the greatest preference for p50 homodimers in comparison to c-Rel or RelA homodimers in prior in vitro PBM experiments.^31^

An analysis of enriched motifs in bins representing the full spectrum of c-Rel/p50 RPKM ratios revealed that all three NF-κB motifs from the Homer program exhibit their lowest enrichment in bins containing c-Rel preferential peaks, most likely reflecting once again the ability of c-Rel homodimers to bind divergent motifs and half-sites. The greatest differences between p50-preferential and c-Rel-preferential peaks were observed with p50 and c-Rel de novo motifs defined above (Figure 5C, de novo p50 and de novo c-Rel motifs).

Notably, a p50 ChIP-seq peak was not observed at the *Il12b* promoter (data not shown). The absence of a p50 peak is consistent with the hypothesis that the *Il12b* promoter is regulated by c-Rel homodimers. However, we cannot exclude the possibility that the absence of a p50 peak is due to relatively poor p50 antibody quality.

To summarize, these ChIP-seq studies provide strong confirmation that the DNA binding characteristics of various NF-κB dimers observed in biochemical experiments reflect dimer-specific binding preferences in vivo at a genome-wide scale. In particular, c-Rel preferential binding occurs at genomic locations that often lack known NF-κB dimeric consensus sequences recognized by heterodimeric species (RelA:p50 and c-Rel:50) or by p50 homodimers. Of greater importance, c-Rel preferential binding in vivo was observed at the *Il12b* promoter and at the promoters of a few other genes that exhibit relatively strong c-Rel-dependence in macrophages, providing a connection between c-Rel-dependent transcriptional induction and the intrinsic binding differences between c-Rel and RelA dimers.

### Late Emergence of the c-Rel-RelA DNA Binding Difference During Vertebrate Evolution

The above results suggest that the duplication and divergence of the genes encoding c-Rel and RelA coincided with the emergence of adaptive immunity and that a primary function of c-Rel is to regulate adaptive immunity. Furthermore, our findings strongly suggest that c-Rel-dependent induction of *Il12b* transcription in macrophages is due at least in part to the strong DNA-binding affinity of c-Rel homodimers for non-consensus motifs in comparison to other NF-κB dimeric species. Together, these results suggest that the initial evolutionary divergence of c-Rel and RelA in early vertebrates may have been driven by mutations that resulted in c-Rel-RelA DNA-binding affinity differences, thereby facilitating neofunctionalization of c-Rel.

To test this hypothesis, we isolated the RHRs of c-Rel and RelA from elephant shark (cartilaginous fish), zebra fish (bony fish), frogs (amphibian), chicken, and humans. The proteins were expressed in HEK 293 cells and extracts prepared. Competition time course experiments were then performed by EMSA to measure relative binding affinities of the two proteins in each species (Figure 6A). For these experiments, the proteins were pre-bound to a radiolabeled oligonucleotide containing a consensus NF-κB motif. Then, a large excess of an unlabeled oligonucleotide containing the same sequence was added and the samples were loaded onto a polyacrylamide gel after different times. When the bound protein dissociates from the radiolabeled oligonucleotide, re-binding will almost always be to the excess unlabeled oligonucleotide, such that the decline is radiolabeled protein-DNA complex provides a measure of protein off-rate and relative binding affinity.

**Figure 6.**
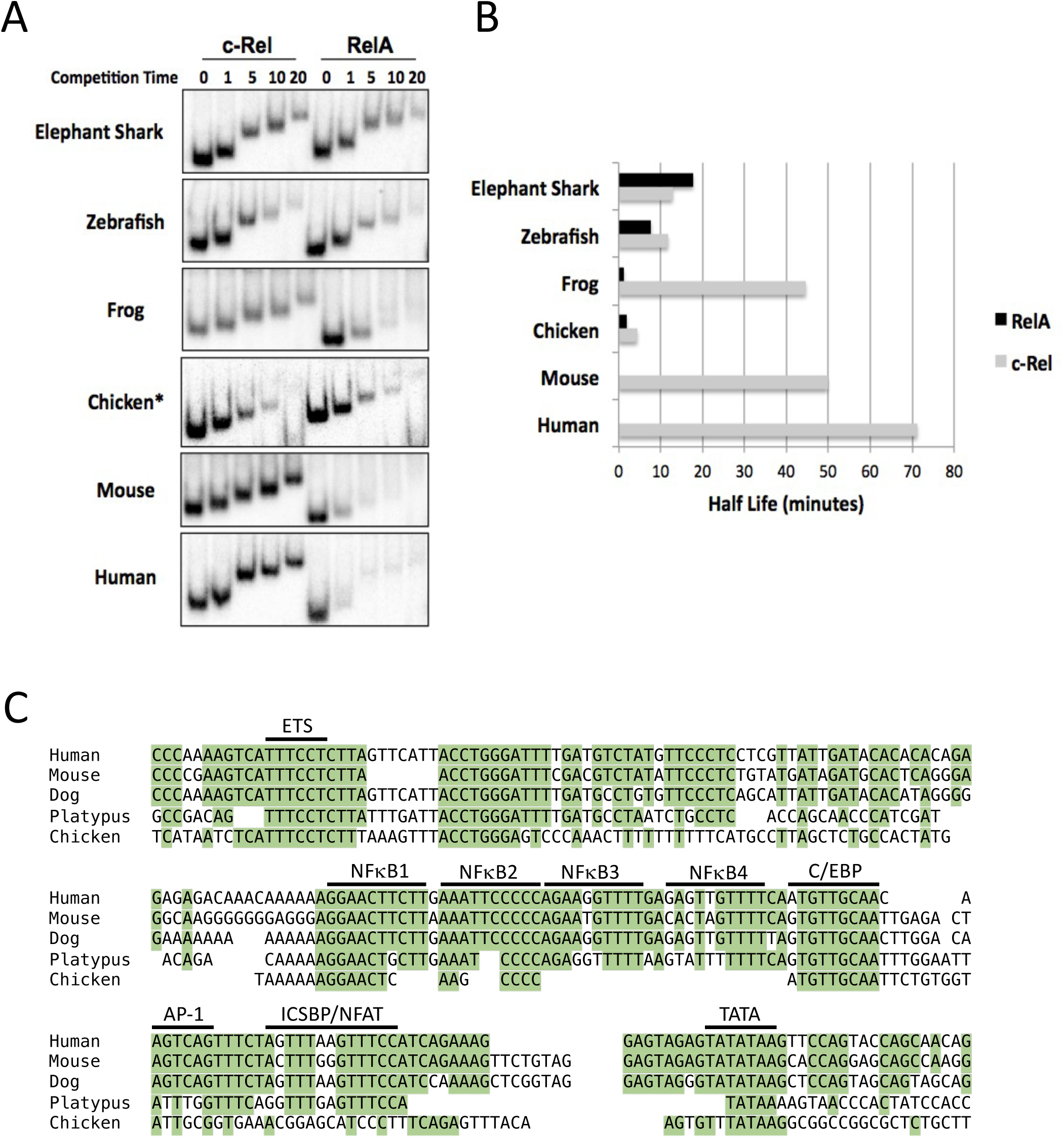
Late Evolution of the c-Rel and RelA DNA-Binding Differences. (A) The RHRs of RelA and c-Rel from six vertebrate species were over-expressed in HEK293T cells. Nuclear extracts were prepared and EMSA off-rate experiments were performed with a radiolabeled, double-stranded oligonucleotide probe containing a consensus NF-κB motif. For these experiments, after pre-binding of the protein to the radiolabeled probe, a large excess of the same oligonucleotide without the radiolabel was added. At 0, 1, 5, 10, and 20 minutes after addition of the unlabeled oligonucleotide, a sample was added to the native polyacrylamide gel. Since protein that releases from the radiolabeled probe is far more likely to re-bind the unlabeled oligonucleotide in excess, measurement of the rate of loss of the protein-DNA complex provides an approximate measure of its half-life. (B) The half-lives of the protein-DNA complexes examined in panel A are shown. The results are representative of at least three independent replicates for each pair of paralogs. (C) The sequences of the *Il12b* promoter from multiple vertebrate species are compared.

Using this approach, the large off-rate difference between mouse c-Rel and RelA was readily observed, as RelA binding to the labeled oligonucleotide declined substantially one-minute after additional of the unlabeled oligonucleotide (Figure 6A,B). In contrast, the c-Rel complex was stable throughout the 20-minute time course. A similar off-rate difference was observed with the human and frog orthologs of c-Rel and RelA. Surprisingly, however, RelA and c-Rel proteins from elephant shark, zebrafish, and chicken exhibited similar off-rates (Figure 6A,B). These results suggest that, in contrast to the above hypothesis, the initial evolutionary divergence of c-Rel and RelA was not driven by the key DNA-binding affinity difference we have defined. Instead, the divergence was likely supported by other differences either in the proteins or their expression patterns (see Discussion). Notably, we also cloned and expressed the Rel cDNA from sea lamprey that exhibits the strongest homology to c-Rel and RelA; by EMSA, this protein bound well to NF-κB consensus sequences in a sequence-specific manner, with an off-rate intermediate between the mammalian c-Rel and RelA off-rates (data not shown).

Additional insights emerged from a comparison of the *Il12b* promoter sequences (Figure 6C). The four potential *Il12b* NF-κB motifs are well-conserved among placental mammals (human, mouse, and dog in Figure 6C), and they are reasonably well-conserved in a marsupial (Figure 6C, platypus). We were also able to align the chicken *Il12b* promoter sequence because the transcription start site and promoter were previously identified.^42^ Although the *Il12b* NF-κB1 and NF-κB2 motifs display significant conservation in chicken, the NF-κB3 and NF-κB4 motifs are absent, with the highly conserved C/EBP motif (previously shown to be critical for mouse *Il12b* promoter function)^43^ located immediately downstream of the chicken NF-κB2 motif. Because of limited homology between promoters from distant species and the absence of mapped transcription start sites, we were unable to identify *Il12b* promoter sequences in amphibian or fish genomes. Nevertheless, the absence of the NF-κB3 and NF-κB4 motifs in the *Il12b* promoter from chicken is consistent with the hypothesis that the divergent DNA-binding properties of c-Rel and RelA may have emerged at a relatively late stage of vertebrate evolution.

### Structural Analysis of the c-Rel versus RelA DNA Affinity Difference

Finally, to identify structural features that may explain the c-Rel-RelA DNA affinity difference, and that may provide insight into the emergence of this affinity difference during vertebrate evolution, we solved the structure of the mouse RelA (C46) chimeric protein homodimer bound to a consensus NF-κB recognition motif. Comparisons between previously reported RelA and c-Rel homodimer co-crystals were of uncertain accuracy because prior structures with the two proteins were solved using different DNA motifs. For our analysis, we used the same consensus recognition sequence (GGGAATTTCC) employed for a prior mouse RelA homodimer structure.^43^ Our focus on the RelA (C46) chimeric protein provided the added benefit of allowing a direct comparison between two proteins that differed only by the 46 residues known to confer the DNA affinity difference.^28,31^

The structure of the RelA (C46) protein-DNA complex was solved by molecular replacement using 2RAM.pdb as the search model and was refined to 3.1Å with an R_free_ of 32.88%. The structures of RelA (C46) (Figure 7A, green) and of RelA bound to the same DNA sequence (cyan)^44^ are very similar. Notably, the structures are also highly similar to that of a c-Rel homodimer (Figure 7A, magenta) bound to a different sequence,^30^ as well as to RelA homodimers bound to different sequences (data now shown).^44,45^ In addition, the relative orientation of the N- and C-terminal subdomains (RHR-N and RHR-C) of one monomer (Figure 7A, monomer A, left) is very similar between the three proteins. However, the orientation of the second monomer (monomer B, right) shows significant differences between the RelA and RelA (C46) complexes in comparison to the c-Rel complex. This difference is likely due to the use of a different DNA sequence for the c-Rel structure, as similar structural variations were observed with RelA bound to different DNA sequences.^44^

**Figure 7.**
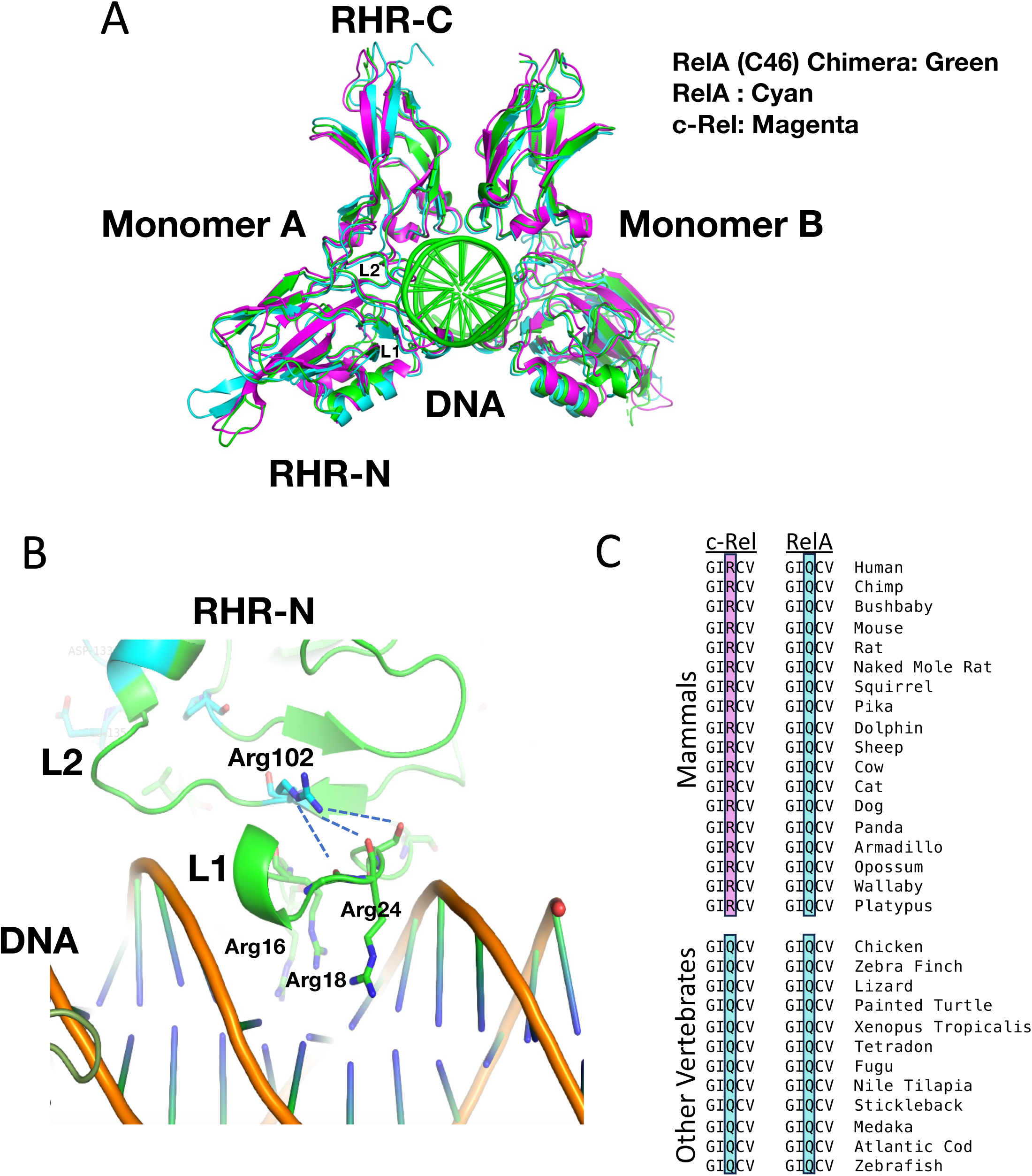
Structure Analysis of RelA Homodimer and RelA (C46) Homodimer Co-Crystals. (A) A comparison is shown between the structure of the mouse RelA (C46) chimeric RHR homodimer (green), a mouse RelA RHR homodimer (cyan), and a mouse c-Rel RHR homodimer (magenta). Figures were prepared with PyMOL (Molecular Graphics System, Version 2.5.6, Schrödinger, LLC). The RelA (C46) and RelA homodimers were bound to the same NF-κB motif sequence (GGGAATTTCC), whereas the c-Rel homodimers were bound to a distinct non-consensus sequence (GGGTTTAAAGAAATTCCAGA).^30^ The N-terminal and C-terminal regions of the RHR (RHR-N and RHR-C, respectively), the two monomers of the homodimer, and the DNA are labeled. (B) Within one RHR monomer, the positioning of the c-Rel Arg102 residue thought to be a major contributor to the enhanced DNA-binding affinity of c-Rel homodimers for DNA is shown. This residue’s position is shown relative to the L1 and L2 loops, and to the DNA-interacting residues Arg16, Arg18, and Arg24. Possibly stabilizing interactions are shown with dashed lines. (C) The residues corresponding to Arg102, as well as two flanking residues on either side, are shown for both RelA and c-Rel for 18 mammalian and 12 non-mammalian vertebrate species. The sequences were derived from the UCSC Genome Browser Comparative Genomics function.

Focusing attention on one monomer, the DNA binding surface structure was also found to be highly similar between the three complexes (Figure S7A). The DNA recognition loop L1, a key DNA binding element of RHR, is positioned in the DNA major groove similarly in all three complexes. The DNA binding loop L2, which interacts with the DNA minor groove, is also similarly oriented in all three complexes. Furthermore, the direct DNA binding surface residues of RelA (C46) are nearly identical to those of RelA (with only a few highly homologs substitutions such as Arg to Lys). These observations suggest that the different DNA binding affinities between RelA (C46) and RelA (also between c-Rel and RelA) are unlikely due to differences in the DNA binding surface. Although many of the residues at the RelA (C46) DNA binding surface show different side-chain conformations from their corresponding counterparts in RelA, these differences may not be responsible for the different DNA bidding activities observed between RelA (C46) and RelA. This is because RelA bound to different DNA sequences^44,45^ showed similar side chain orientation variations (data not shown).

We highlighted c-Rel specific residues (stick model, cyan) on the structure of the RelA (C46) chimera (Figure S7B, only the RNR-N of one monomer and its bound DNA are shown). None of the c-Rel-specific residues contact DNA directly. We next grouped the c-Rel specific residues into three clusters. Residues in cluster I are located in the loop regions far away from the protein core and the DNA binding surfaces. These residues may have little impact on DNA binding but may be involved in interactions with other proteins. Residues in cluster II are involved in the packing interaction with the beta-sandwich core of the proteins; this region is also connected with the DNA binding loop L2 that binds DNA in the minor groove. These c-Rel specific residues may therefore impact DNA binding by modulating the conformation flexibility of the protein fold and conformation flexibility of the DNA binding loop L2.

Interestingly, the c-Rel-specific residue that is most likely to have a major impact on DNA binding affinity is Arg102, a sole residue in cluster III. As shown in Figure 7B, the positively charged Arg102 is located behind L1 loop and its long side chain and the guanidinium head form a network of hydrogen bonding interactions (dashed lines) with the backbone carbonyl groups on the L1 loop. This interaction can stabilize the L1 loop and its three arginine residues (Arg16, Arg18, and Arg 24) in a favorable conformation for binding to DNA. In RelA, the corresponding residue is a glutamine (Q), which is neutral and shorter and could not form the same L1 stabilization interactions as Arg102 in c-Rel. Notably, although Arg102 is likely to make a major contribution to the enhanced DNA-binding affinity of c-Rel, it must act in concert with other residues that differ between c-Rel and RelA (such as those in cluster II in Figure S7B) because, in our extensive efforts to localize residues responsible for the c-Rel-RelA DNA affinity difference,^28^ the difference was observed only when the entire 46-residue c-Rel region was introduced into RelA.

Most importantly, an analysis of the c-Rel and RelA sequences in diverse species revealed that Arg102 is perfectly conserved in the c-Rel orthologues of all 41 mammalian species available in the UCSC Genome Browser Comparative Genomics function (Figures 7C, S1 and data not shown), Similarly, the corresponding RelA residue, Gln102, is perfectly conserved in all of the mammalian species (Figures 7C, S1 and data not shown). However, in all non-mammalian vertebrate species in the comparative sequence analysis, this residue is a glutamine in both the c-Rel and RelA orthologues (Figures 7C, S1 and data not shown). This finding supports the notion that the Gln102 to Arg102 change in a mammalian ancestor contributed to the evolution of the c-Rel-RelA affinity difference. Notably, although a c-Rel-RelA DNA affinity difference was also observed with frog (*X. laevis*) proteins in Figure 6A, the *X. laevis* c-Rel has a glutamine rather than an arginine at position 102. This observation suggests that, in *X. laevis*, other sequence changes may have led to the c-Rel-RelA DNA affinity difference. *X. laevis* is unusual, however, in having two apparent c-Rel orthologs within its genome (data not shown), only one of which we have examined biochemically.

## DISCUSSION

We combined biochemical, genomic, functional, structural, and evolutionary analyses to examine a critical difference between the closely related NF-κB paralogs, c-Rel and RelA. Our results provide a clear examine of the acquisition of a structural change in a transcription factor family member at a late stage of vertebrate evolution, resulting in a greatly increased intrinsic DNA-binding affinity with major functional consequences. PBM and SPR experiments revealed previously unrecognizable motifs that can be bound by c-Rel homodimers due to their intrinsic ability to bind DNA with an enhanced affinity in comparison to RelA homodimers. Biochemical and functional analyses showed that c-Rel homodimers can bind selectively to four tandem motifs in the promoter of the *Il12b* gene; two of these motifs would have been unrecognizeable prior to the PBM analysis. ChIP-seq experiments demonstrated that the distinct binding properties of c-Rel, RelA, and p50 dimers defined biochemically can largely explain their preferential distributions across the genome. Moreover, despite the dramatic c-Rel-RelA binding differences and the strong evidence of functional importance, the distinct binding properties were not observed with c-Rel and RelA orthologs from early gnathostomes, suggesting that this key difference arose at a relatively late stage of gnathostome evolution. Finally, structural studies identified a key mammal-specific c-Rel residue that is likely to contribute to this late-evolving property of c-Rel.

The absence of intrinsic DNA-binding differences between RelA and c-Rel in early gnathostomes raises the question of what other properties of the two proteins and their genes drove their divergence and specialization. One possible contributor to their specialization is their differential expression. Differential expression of the *Rela* and *Rel* genes is well-known,^19^ with *Rela* expressed at a high level much more broadly across mammalian tissues than *Rel*. Differences in the regulation of stimulus-responsive nuclear translocation of RelA and c-Rel complexes have also been described in both innate and adaptive immune cell types.^46–49^ These expression differences may have allowed expansion of NF-κB’s regulatory potential.

Differences in the RelA and c-Rel C-terminal transactivation domains may have also contributed to the divergence and specialization of the two family members, possibly contributing to their regulation of distinct sets of genes in different settings. The transactivation domains of RelA and c-Rel are poorly conserved through evolution and our prior chimeric protein studies showed that the C-terminal transactivation domains of RelA and c-Rel are interchangeable for the activation of the endogenous *Il12b* gene in *Rel^-/-^* macrophages. Nevertheless, the transactivation domain of mammalian RelA supports a well-documented interaction with the p300/CBP co-activators^15,50^ and this interaction does not appear to be supported by c-Rel’s transactivation domain.^25^ The p300/CBP interaction of RelA has been shown to be critical for the activation of a distinct subset of NF-κB target genes,^51^ raising the possibility that this c-Rel-RelA difference may have contributed to the initial divergence of the two genes.

The late evolution of the DNA binding difference between RelA and c-Rel suggests that this intrinsic biochemical difference may support immunoregulatory functions that were not important in early gnathostomes. Alternatively, conserved immunoregulatory functions of c-Rel that did not rely on the DNA affinity difference in early gnathostomes came to rely on this affinity difference in later species.

Consistent with the possibility that the distinct DNA-binding properties of c-Rel and RelA may have been important for the evolution of advanced tolerance mechanisms, c-Rel and RelA both play critical but distinct roles in the development and function of Tregs,^52–54^ which are known to have experienced changes in their regulatory strategies at late stages of vertebrate evolution.^7^ These late changes in Treg regulation have been hypothesized to support fetal tolerance.^7^ Notably, the Arg102 residue that is likely to be a key contributor to the c-Rel-RelA affinity difference is observed in c-Rel from both placental and non-placental mammals, suggesting that its emergence does not precisely coincide with the need for placental tolerance.

Whether the DNA-binding differences described here are critical for the distinct activities of RelA and c-Rel in Treg development and function remains to be determined.^53,54^ It also is not yet known whether the binding differences are important for the distinct functions that have been documented for RelA and c-RelA in other cells of the adaptive immune system. In support of this possibility, the first DNA motif proposed to confer c-Rel-selectivity in T cells is located in a control region for the *Il2* gene, referred to as the CD28 response element.^55^ Although both RelA- and c-Rel-containing dimers can bind this motif,^56,57^ it possesses a non-consensus sequence that would be predicted to allow preferential binding and activation by c-Rel homodimers.

Another unanswered question is why *Il12b* acquired a critical requirement for c-Rel for its activation in macrophages, whereas many other putative NF-κB target genes appear to be activated redundantly by RelA and c-Rel complexes. This finding suggests that *Il12b* would not function properly if it were regulated similarly to other NF-κB target genes, thereby mandating its requirement for c-Rel. A likely reason for the emergence of a c-Rel requirement would be because c-Rel dimers are subject to unique regulatory mechanisms. Although dimers containing c-Rel and RelA are thought to be maintained in an inactive state by the same IκBα and IκBβ proteins, activation differences have occasionally been reported, including unique requirements for Caspase 8^49^ or cellular nucleic acid binding protein (CNAP)^58^ for c-Rel activation in some settings. Proper regulation of *Il12b* expression by c-Rel may rely on these or other unique mechanisms.

RelA and c-Rel expression kinetics are also regulated differently in activated macrophages, with c-Rel expression more prolonged than RelA expression in response to some stimuli. This difference can allow c-Rel dimers to play a broader role in inflammatory gene regulation after RelA dimers are downregulated. These c-Rel-RelA expression differences would allow a broad range of inflammatory genes to be sensitive to the modulation of c-Rel activity, as recently described for the regulation of c-Rel activity by very long chain ceramides associated with anti-inflammatory pathways.^59^

Finally, the absence of the intrinsic DNA-binding difference between RelA and c-Rel homodimers in early vertebrates raises the question of whether *Il12b* induction in early vertebrates remains c-Rel-dependent albeit with the c-Rel dependence due to a different mechanism, or whether *Il12b* induction in these species is perhaps regulated by c-Rel-independent mechanisms. Possibly related to this question, *Il12b* activation in mammals is known to be c-Rel independent in some settings.^60^ This knowledge adds strength to the possibility that *Il12b* activation in early vertebrates may rely entirely on c-Rel-independent mechanisms that remain intact in mammals but have been enhanced by a critical c-Rel-dependent regulatory mechanism in key cell types and settings.

## EXPERIMENTAL PROCEDURES

### Phylogenetic Analysis and Expression of NF-κB cDNAs from Diverse Species

The unrooted phylogenetic tree was constructed with BEAST 1.0 software^61^ using approximately 200 residues of the RHRs shown in Figure S1 and corresponding regions from the other proteins shown in Figure 1A.

RHR cDNAs were amplified from mRNA or cDNA libraries by PCR and cloned into the pcDNA3 vector with an N-terminal Flag epitope tag. c-Rel and RelA RHR cDNAs were from mouse^28^, human, chicken, frog (*Xenopus laevis*), zebrafish, and elephant shark. The sea lamprey Rel1 cDNA was cloned from mRNA isolated from freshly isolated blood. The recombinant proteins were expressed in HEK 293T cells following transient transfection of the expression plasmids. Nuclear extracts were prepared for EMSA as previously described.^28,43^

### X-Ray Crystallography

The RelA(C46) RHR chimeric protein was expressed and purified as described.^31^ The protein was crystallized at 18°C using the hanging-drop, vapor-diffusion method with a reservoir solution of 0.1 M Bis-Tris propane (pH 6.5), 0.2 M sodium/potassium phosphate, 20% PEG 3350. 10 mM DTT and 0.5% BOG (beta octyl glucoside). X-ray data were collected at ALS beamline 8.2.2. Data were processed and scaled with the HKL2000 software package.^62^ Initial phase information was obtained by molecular replacement method with Phaser^63^ using the crystal structure of NF-κB p65(PDB 2RAM) as the search model. The structural model was refined using PHENIX^64^ and modified with COOT.^65^ Data collection, phasing and refinement statistics are summarized in Figure S8. PyMOL (Molecular Graphics System, Version 2.5.6, Schrödinger, LLC) was used to generate the figures. The structural information has been deposited in the Protein Data Bank (8U9L.pdb; https://doi.org/10.2210/pdb8U9L/pdb). Comparisons are shown to RelA homodimers bound to the same DNA motif^44^ (Chen et al., 2000) and to c-Rel homodimers bound to a different DNA sequence (1GJI.pdb).^30^

### PBM and SPR Experiments

The PBM experiments were described previously,^31^ with further computational scrutiny of bound oligonucleotides revealing frequent binding to oligonucleotides containing the tandem T:A bps. SPR was performed as described,^31^ using purified recombinant RHRs as described.^31^

### Transient Transfection Experiments

The mouse *Il12b* promoter (−355 to +55) was cloned into the pGL4.10 vector (Promega). Motif mutations were introduced using the GENEART site-directed mutagenesis system (Invitrogen). C-terminal single Flag tag versions of full-length cDNAs encoding mouse RelA, c-Rel, and p50 were cloned into pcDNA3 vectors. 293T cells, grown in DMEM with 10%FBS, were transfected with Lipofectamine 2000 (Invitrogen) in 24-well plates. 1-8,000 ng of the expression plasmid were co-transfected with 20ng of the *Il12b*-pGL4 vectors. 1 ng of a TK-Renilla luciferase vector was included as a transfection control. Cells were collected 24-hr post transfection and firefly and renilla luciferase were analyzed with the Dual Luciferase Reporter Assay system (Promega). Normalization was performed using Western blot analysis for protein concentration, renilla luciferase for transfection efficiency, and empty vector firefly/renilla values for background signal.

### Engineering and Analysis of Mutant *Il12b-Gfp* BACs

Recombination in E. coli by standard procedures^66^ was used to introduce a *Gfp* cDNA into the second exon of a 200 kb BAC spanning the mouse *Il12b* locus (Doty et al., unpublished results). Recombination in E. coli was then used to introduce substitution mutations into *Il12b* promoter motifs (see Figure S3B). The recombinant WT and mutant *Il12b-Gfp* BACs were introduced into a feeder-free mouse ESC line. Clonal lines were expanded and single-copy integrants were identified by qPCR and Southern blot analysis (Figure S3 and data not shown).

BAC-containing ESC were differentiated into macrophages as described,^67^ with confirmation of differentiation into myeloid progenitors and macrophages confirmed by flow cytometry (Figure S3A). Macrophages were then stimulated with lipid A (100ng/ml) for 2 hr, following by qRT-PCR analysis of BAC-derived *Gfp* mRNA. Endogenous *Il12b* mRNA was monitored in each line by qRT-PCR as a positive control.

### Electrophoretic Mobility Shift Assays (EMSAs) and Immunoblot Assays

Purified proteins (Figures 4 and S1-S4) or nuclear extracts from transfected HEK293T cells (Figure 7) were incubated with radiolabeled probe (in 25μL total volume) in the presence of 10mM Tris-HCl, 150mM NaCl, 1mM DTT, 1mM EDTA, 5% glycerol, 80ng/μL dI-dC, and 200ng/μL BSA, with limited amounts of double-stranded ^32^P-labeled probes (∼10^-11^ M). Each reaction was incubated at 4°C for 30 minutes before gel electrophoresis. Gel shift assays were run as described.^43^ Band intensities were quantified using ImageQuant (GE).

Immunoblots were performed as previously described,^31^ using antibodies directed against RelA or c-Rel.

### Nascent Transcript RNA-seq

Bone marrow-derived macrophages were prepared from C57BL/6 and *Rel^-/-^*mice as described.^68^ Macrophages were activated on day 6 with lipid A for the time indicated in each experiment (Sigma-Aldrich). Nascent transcript RNA-seq was performed and the data were analyzed as described.^68,69^

### ChIP-Seq Analysis

ChIP-seq was performed as described^68,70^ with anti-RelA, anti-c-Rel, and anti-p50 antibodies (Cell Signaling, Inc. 8242, 68489, and 13586, respectively). Approximately 10 million BMDMs were used per sample from C57BL/6 mice aged 8-12 weeks. After crosslinking with 1mM DSG and 1% PFA, cells were sonicated on a Covaris M220 focused ultrasonicator to 200-500-bp DNA fragments. ChIP-seq libraries were prepared and peaks were called as described.^68,71^ Read were aligned using Hisat2 to the NCIB37/mm9 mouse genome. To compare peaks across multiple samples, a master probe was generated with BED Tools.^72^

## ACKNOWLEDGMENTS

We thank David Schatz for critical reading of the manuscript. This work was supported by NIH grants R01GM086372, R01AI073868, R01CA127279, and P50AR063020 (to S.T.S.), F31AI157267 (to A.E.D.), T32CA009120 (to A.B.C. and K.J.W.), and T32AI007323 (to A.E.D. and S.D.P.) S.T.S. is a Senior Scientific Officer of the Howard Hughes Medical Institute.

## SUPPLEMENTAL FIGURE LEGENDS

**Figure S1.**
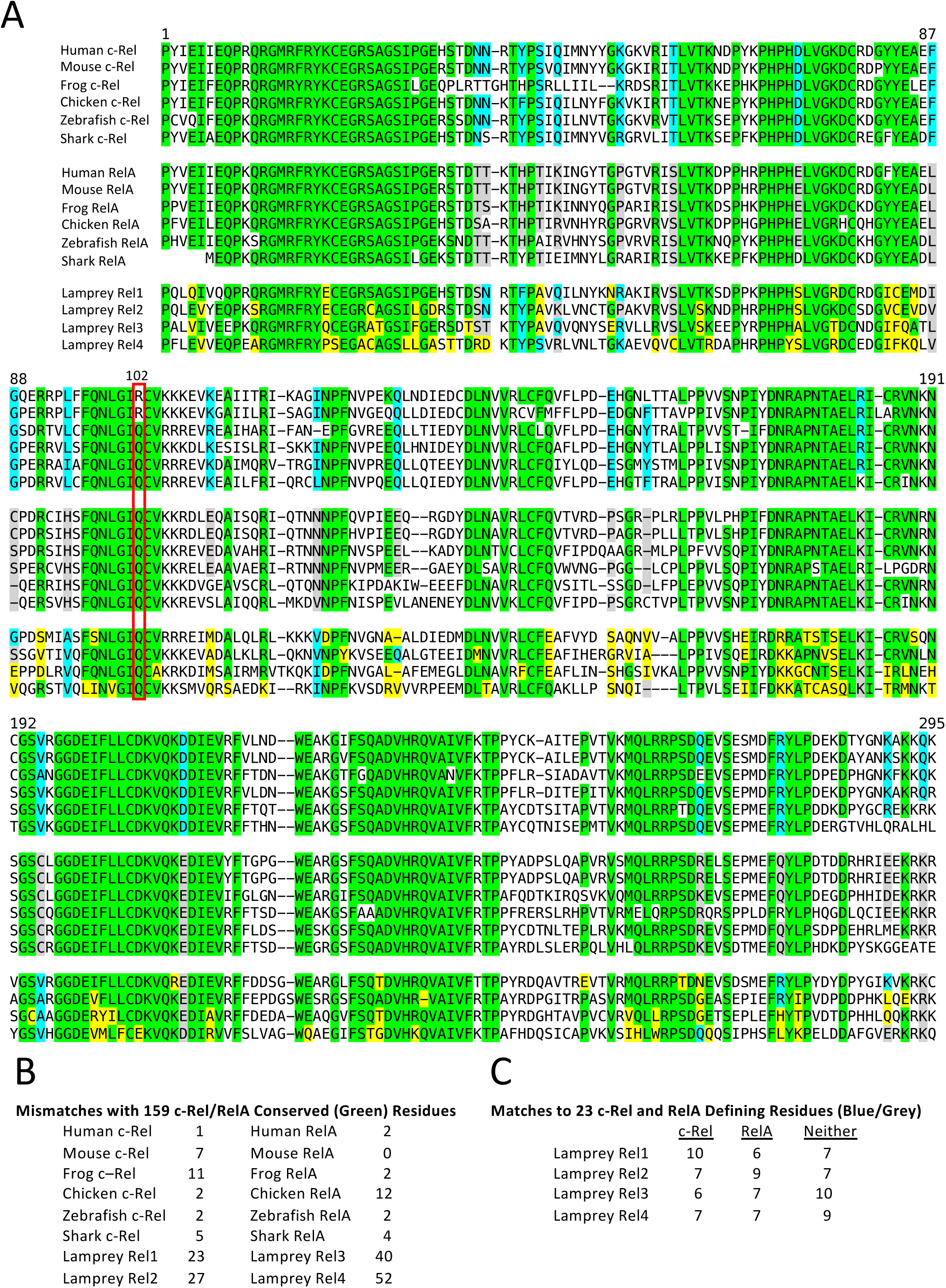
Comparison of c-Rel and RelA RHR Sequences from Six Gnathastome Species to Four Rel RHRs from Sea Lamprey. (A) RHR sequences are shown for c-Rel and RelA orthologs from six gnathostome species and from four predicted Rel proteins identified in the sea lamprey genome. The fifth sea lamprey RHR is not shown due to its extensive homology to the NF-κB p50 and p52 RHRs. Amino acids are shaded in green that are perfectly conversed in at least five of the six species for c-Rel and in at least five of the six species for RelA. These amino acids are also in green in the lamprey RHRs if they match the gnathostome residue, and it yellow if they do not match. 23 “defining” residues that most consistently distinguish the c-Rel RHRs from the RelA RHRs are highlighted in blue (c-Rel) and grey (RelA). These 23 residues are sometimes identical in all six gnathostome c-Rel or RelA RHRs, or they may be members of the same amino acid class. These residues are shaded in blue or grey in the lamprey RHR sequences if they match the characteristics of the c-Rel or RelA defining residues, respectively, and they are shaded in yellow if they do not match either the c-Rel or RelA characteristics. Arg102/Gln102, suggested by X-ray crystallography to be critical for the c-Rel-RelA DNA affinity difference is highlighted with a red box. (B) For the 159 residues shaded in green in panel A, the number of mismatches in each of the sixteen RHRs is shown. Note that all four of the lamprey RHR contain many more mismatches than any of the RHRs in the six gnathostome species. (C) For the 23 defining residues that most consistently distinguish the c-Rel RHRs from the RelA RHRs, the numbers of residues in each lamprey RHR that match the c-Rel residue, match the RelA residue, or do not match either residue are indicated.

**Figure S2.**
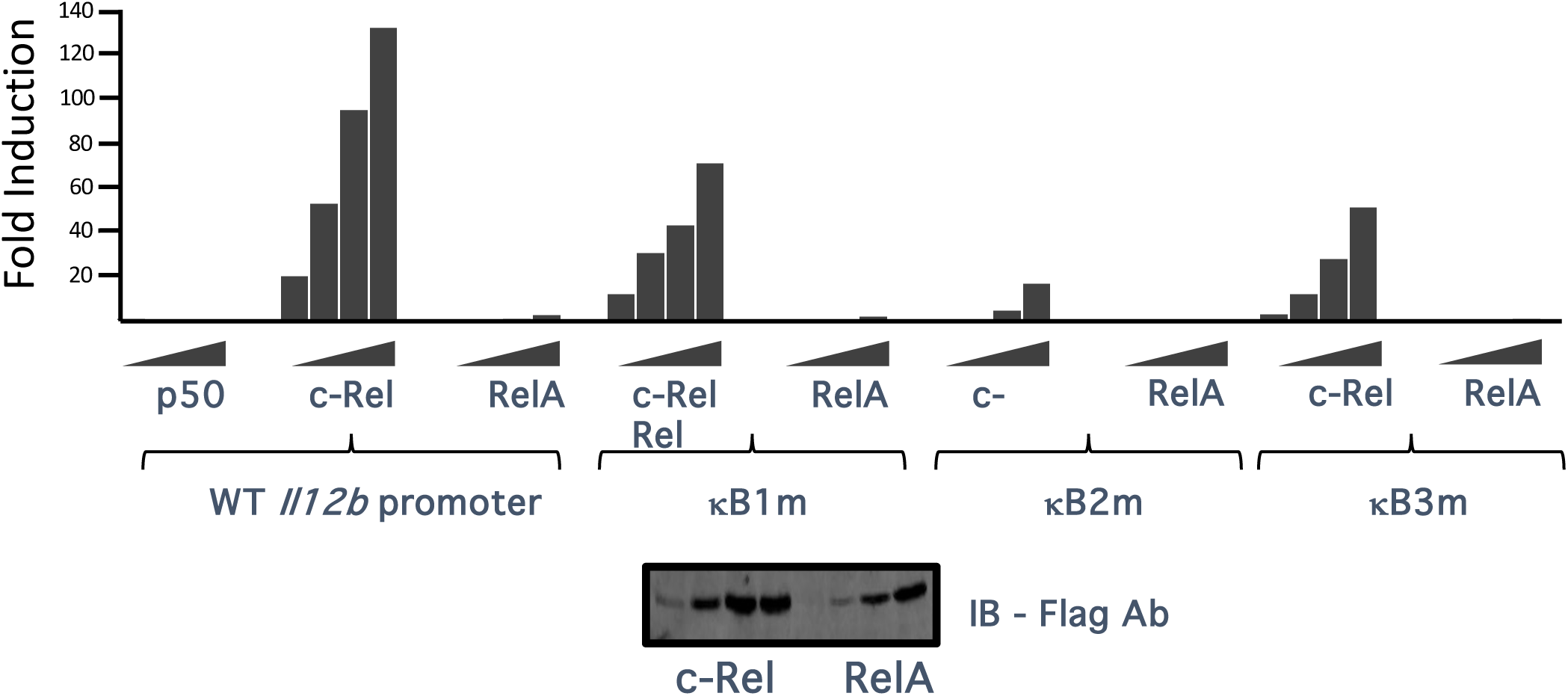
Functional Analysis of Non-Consensus NF-κB Motifs in the Mouse *Il12b* Promoter by c-Rel and RelA Overexpression in HEK 293T Cells. HEK 293T cells were transiently transfected with a mouse *Il12b* promoter-luciferase reporter plasmid and expression plasmids for full-length Flag epitope-tagged RelA, c-Rel, and p50. Luciferase activity was monitored with plasmids containing the WT *Il12b* promoter and plasmids containing substitution mutations in the *Il12b* NF-κB1, NF-κB2, and NF-κB3 motifs. As previously observed (Murphy et al., 1005), c-Rel overexpression activated *Il12b* promoter activity more strongly than RelA overexpression. Promoter activity was compromised to variable extents by mutation of each of the three motifs.

**Figure S3.**
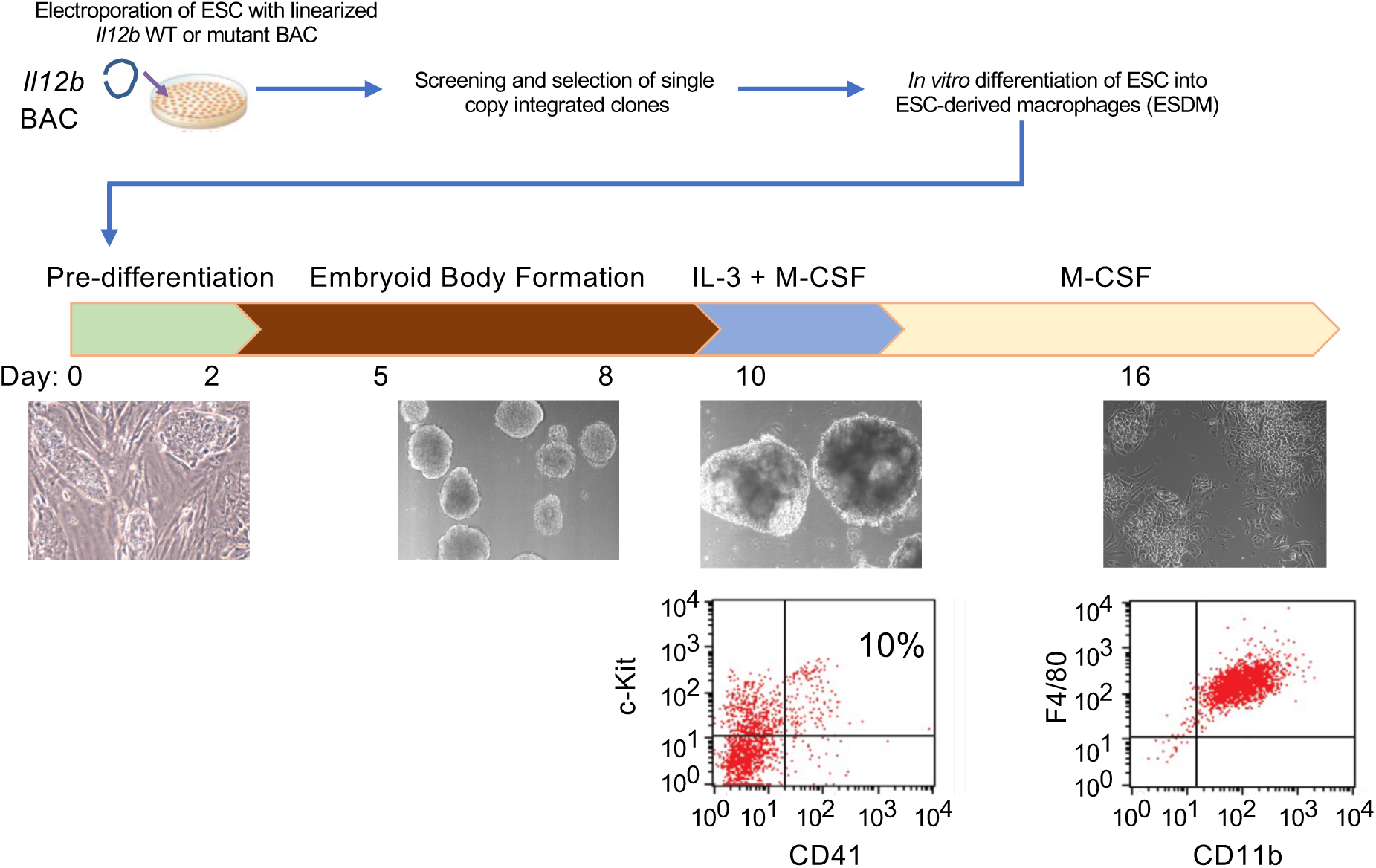
Functional Analysis of Non-Consensus NF-κB Motifs in the Mouse *Il12b* Promoter Using Recombinant BACs in ESC-Derived Macrophages. (A) The strategy used to analyze substitution mutations in the mouse *Il12b* promoter is shown (performed prior to the emergence of CRISPR-Cas9 in mammalian cells). A bacterial artificial chromosome (BAC) spanning the mouse *Il12b* locus was modified by homologous recombination in E. coli to add a *Gfp* cDNA into the *Il12b* transcription unit. This *Il12b-Gfp* BAC was then further modified by homologous recombination in E. coli to introduce mutations into five locations in the *Il12b* promoter: the NF-κB1, NF-κB2, NF-κB3, and NFAT motifs, and a fifth sequence that is not conserved between species. The recombinant BACs were introduced into mouse ESC by electroporation and multiple independent clones for each BAC were isolated and expanded. Following the identification of clones containing single-copy BAC integrants, those clones were expanded and differentiated in vitro into post-mitotic macrophages. The ESC-derived macrophages were then stimulated for 2 hour with LPS. Flow cytometry was used to monitor CD41 and c-kit expression to confirm partial differentiation of ESC into macrophage progenitors. CD11b and F4/80 were then monitored by flow cytometry, demonstrating efficient differentiation into macrophages.

**Figure S4.**
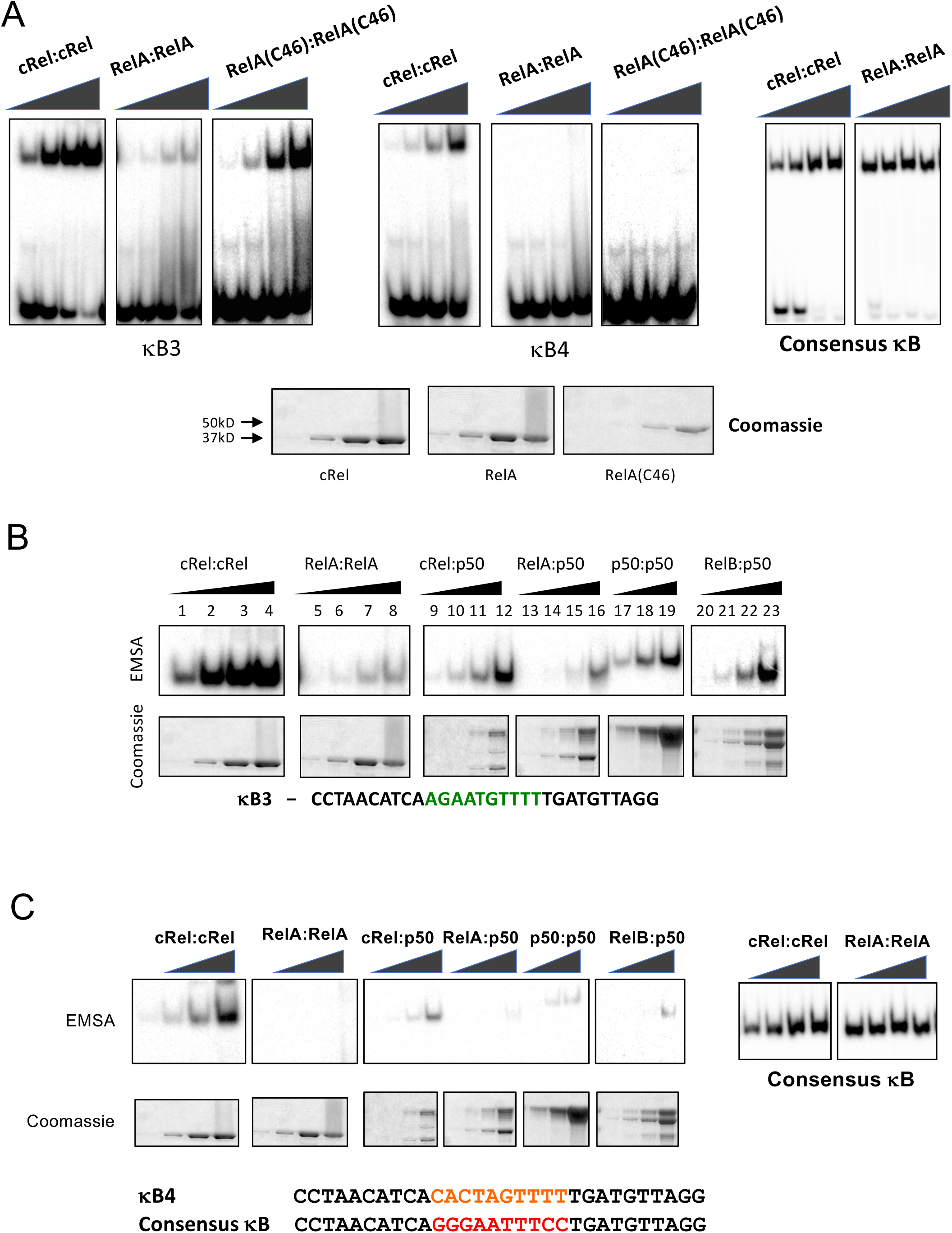
Intrinsic Binding of NF-κB Dimers to the *Il12b* NF-κB3 and NF-κB4 Motifs. (A) EMSAs were performed with recombinant NF-κB RHR dimers expressed in E. coli. Binding was compared with increasing concentrations of c-Rel, RelA, and RelA(C46) homodimers to radiolabeled oligonucleotide problems containing the *Il12b* NF-κB3 and NF-κB4 motifs. Comparable binding of c-Rel and RelA homodimers to a probe containing a consensus NF-κB motif was examined as a control (right). Coomassie stains of the purified proteins are shown at the bottom. (B) EMSAs were performed to examine relative binding strength of various NF-κB RHR homodimers and heterodimers to a radiolabeled probe containing the *Il12b* NF-κB3 sequence. For heterodimers, two NF-κB family members were co-expressed in E. coli. Coomassie stains of the purified are shown at the bottom. (C) EMSAs were performed to examine relative binding strength of various NF-κB RHR homodimers and heterodimers to a radiolabeled probe containing the *Il12b* NF-κB4 sequence. For heterodimers, two NF-κB family members were co-expressed in E. coli. Coomassie stains of the purified are shown at the bottom. Binding of c-Rel and RelA homodimers to a probe containing a consensus NF-κB motif is shown at the right as a control.

**Figure S5.**
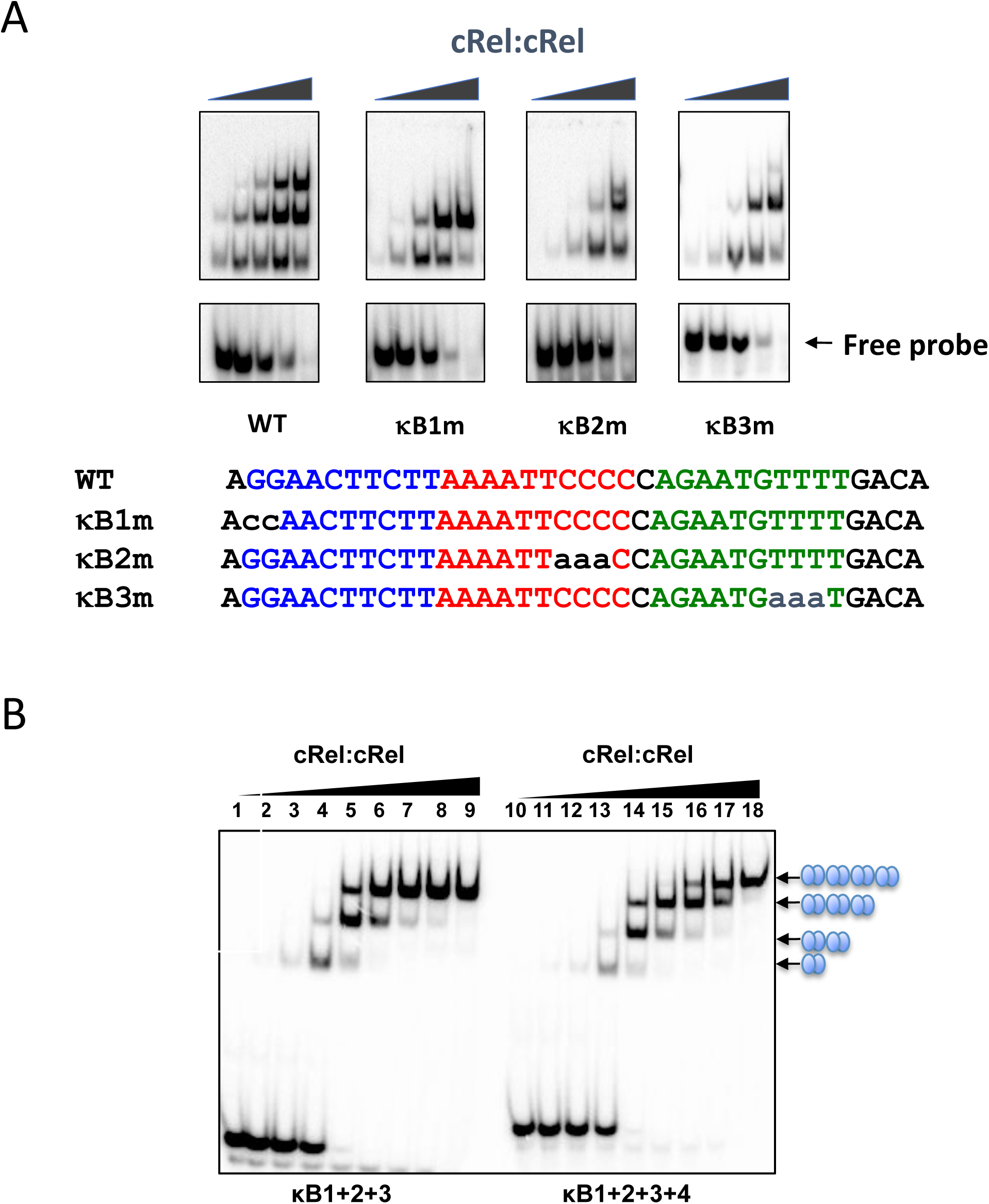
Analysis of c-Rel Binding to *Il12b* Promoter Probes containing Tandem Motifs. (A) EMSAs were used to examine c-Rel RHR binding to probes containing substitution mutations in the *Il12b* NF-κB1, 2, and 3 motifs in a radiolabeled probe containing these three motifs in tandem, as in the WT *Il12b* promoter. (B) An EMSA was used to examine binding of the c-Rel RHR to a radiolabeled probe containing all four tandem *Il12b* NF-κB motifs. A probe containing the *Il12b* NF-κB1 to NF-κB3 motifs is on the left. A probe containing the *Il12b* NF-κB1 to NF-κB4 motifs is on the right.

**Figure S6.**
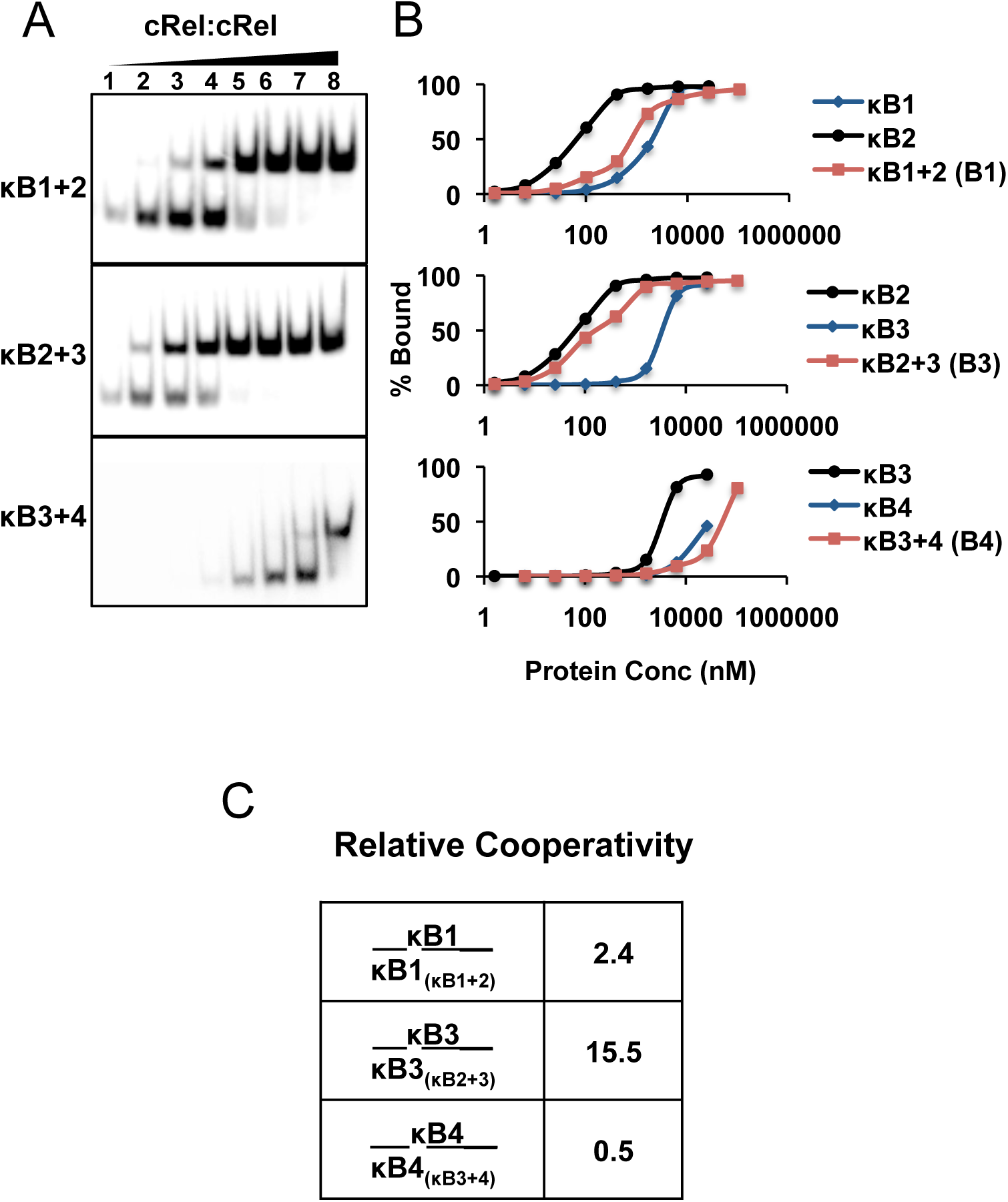
Analysis of Cooperative Binding of c-Rel to the *Il12b* NF-κB Motifs. (A) To examine the possibility of cooperative binding, EMSAs were performed with the c-Rel RHR and probes containing different pairs of *Il12b* NF-κB motifs at their native spacing. For these experiments, increasing concentrations of c-Rel were examined. The percentage of probe that assembles into a complex with the two sites simultaneously was monitored and compared to parallel experiments using probes containing each of the two motifs alone (not shown). (An examination of the complex containing only one bound dimer in the images shown provides a separate measure of binding strength to the highest affinity motif of the two motifs present in each probe.) (B) Quantitative analysis of the data in panel A allows calculation of an approximate cooperativity index. For these calculations, the concentration of protein needed to occupy the lower affinity site on 50% of the probe molecules containing that site alone was divided by the concentration needed to occupy both sites simultaneously in the probe containing both sites. The results show that the presence of the *Il12b* NF-κB2 sequence allows greatly enhanced binding to the lower affinity NF-κB3 sequence. However, compelling cooperativity was not observed with the other motif pairs.

**Figure S7.**
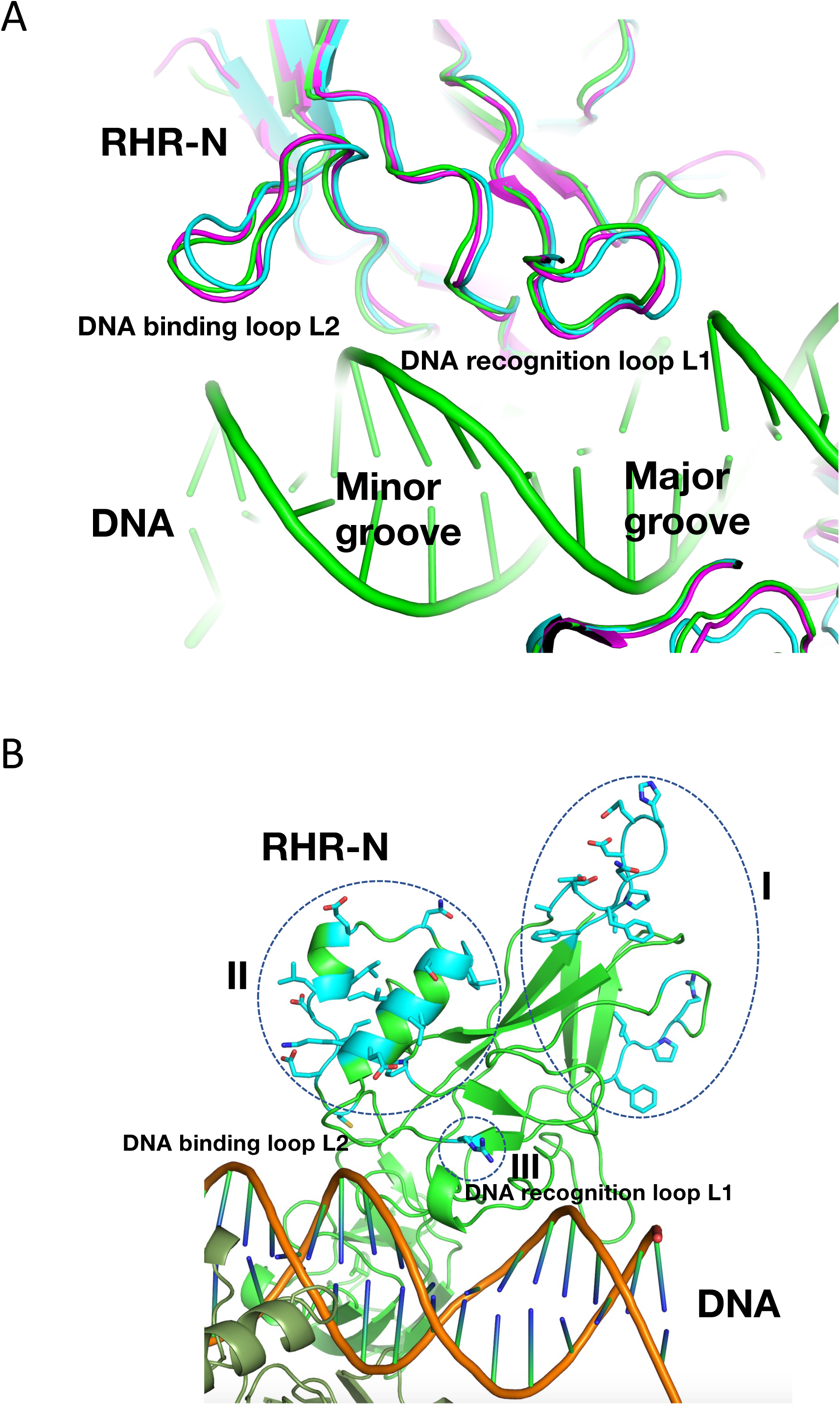
Additional Structure Insights into the c-Rel-RelA DNA Affinity Difference. (A) The extensive similarity between the protein-DNA interfaces observed with the RelA (C46) (green), RelA (cyan), and c-Rel (magenta) proteins is shown. (B) The locations of three clusters of c-Rel-specific RHR residues are highlighted with dashed circles and are labeled (I, II, and III).

**Figure S8.**
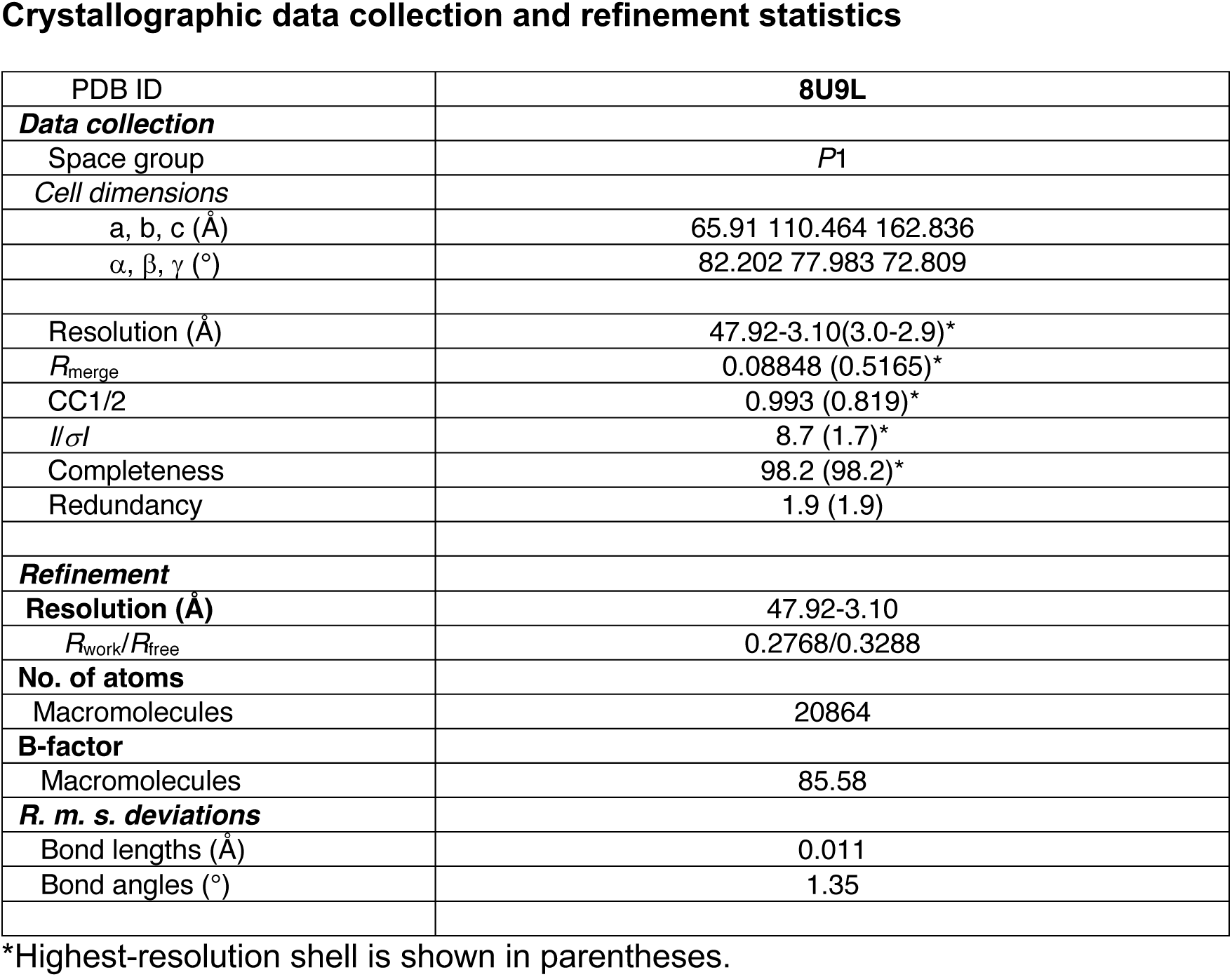
Crystallographic Data Collection and Refinement Statistics. The crystallographic data collection and refinement statistics for the mouse RelA (C46) homodimer-DNA co-crystal analysis are shown.

